# Lateral Axonal Modulation is Required for Stimulus-Specific Olfactory Conditioning in *Drosophila*

**DOI:** 10.1101/2022.06.02.494466

**Authors:** Julia E. Manoim, Andrew M. Davidson, Shirley Weiss, Toshihide Hige, Moshe Parnas

## Abstract

Effective and stimulus-specific learning is essential for animals’ survival. Two major mechanisms are known to aid stimulus-specificity of associative learning. One is accurate stimulus-specific representations in neurons. The second is limited effective temporal window for the reinforcing signals to induce neuromodulation only after sensory stimuli. However, these mechanisms are often imperfect in preventing unspecific associations; different sensory stimuli can be represented by overlapping populations of neurons, and more importantly the reinforcing signals alone can induce neuromodulation even without coincident sensory-evoked neuronal activity. Here, we report a crucial neuromodulatory mechanism that counteracts both limitations and is thereby essential for stimulus specificity of learning. In *Drosophila*, olfactory signals are sparsely represented by cholinergic Kenyon cells (KCs), which receive dopaminergic reinforcing input. We find that KCs have numerous axo-axonic connections mediated by the muscarinic type-B receptor (mAChR-B). By using functional imaging and optogenetic approaches, we show that these axo-axonic connections suppress both odor-evoked calcium responses and dopamine-evoked cAMP signals in neighboring KCs. Strikingly, behavior experiments demonstrate that mAChR-B knockdown in KCs impairs olfactory learning by inducing undesired changes to the valence of an odor that was not associated with the reinforcer. Thus, this local neuromodulation acts in concert with sparse sensory representations and global dopaminergic modulation to achieve effective and accurate memory formation.

**Highlights:**

- Lateral KC axo-axonic connections are mediated by muscarinic type-B receptor
- KC connections suppress odor-evoked calcium responses and dopamine-evoked cAMP
- knockdown of the muscarinic type-B receptor impairs olfactory learning
- Impaired learning is due to changes to the valence of the unconditioned odor

## Introduction

An animal’s survival critically depends on its capacity for sensory discrimination, which enables stimulus-specific modulation of behavior. In associative learning, a particular sensory input is associated with reward or punishment. Across animal phyla, dopamine (DA) plays a central role in mediating those reinforcement signals and inducing synaptic plasticity^1^. The majority of DA signaling is mediated by volume transmission^2^, but this only provides coarse spatial specificity. Neuromodulation must be further restricted to the few synapses whose activity represents the specific sensory stimulus that is being associated with the reinforcer. One important contributing mechanism to such synapse specificity is the requirement of temporal coincidence of DA input and synaptic activity for plasticity. That is, plasticity only occurs at synapses that are co-active with (or were active immediately before) DA input^3–5^. Another important mechanism observed in many cortical areas to help synapse specificity is the sparse coding of sensory representations^6–10^, which minimizes overlap between neuron subpopulations whose activity represents distinct stimuli. This combination of a short eligibility time window and segregated sensory representation is assumed to underlie stimulus specificity of learning^11,12^. However, it is known that as sensory signals ascend to higher brain areas, there is an inherent increase in trial-to-trial variability^13–15^ that might risk the reliability of sparse representation as well as stimulus specificity of learning. In this study, we identified additional novel processes acting locally at axons to enhance synapse specificity of plasticity induction beyond that which can be accomplished at the neuronal population level.

In *Drosophila* olfactory learning, convergence of sensory and reinforcement signals takes place at axons of Kenyon cells (KCs), which are the third-order olfactory neurons constituting the principal neurons of the mushroom body (MB)^16^. Specific DA neurons (DANs) densely innervate specific segments of the KC axon bundles^17–19^. The spatial arrangement of DAN-KC synapses revealed in the EM connectome suggests that DA modulation occurs by volume, rather than local, transmission^20^. Thus, like in vertebrate brains, release of DA is unlikely to be target specific in the MB. Associative learning in flies almost completely depends on G_s_-coupled DA receptor, Dop1R1, expressed in KCs^21,22^. Since one of the classical learning mutant genes identified by genetic screen was *rutabaga* (*rut*), which encodes Ca^2+^-dependent adenylate cyclase abundantly expressed in KCs^23–25^, it is believed that coincidence of odor-evoked Ca^2+^ influx in KCs and DA input triggers cyclic adenosine monophosphate (cAMP)-dependent plasticity^26^. Indeed, KC activation and DA application can synergistically elevate cAMP level in KC axons in a *rut*-dependent manner^27^. Furthermore, simultaneous activation of KCs and DANs can induce robust long-term depression (LTD) at KC to MB output neuron (MBON) synapses, and the plasticity induction strictly depends on the temporal sequence of KC and DAN activity^5,28–30^. Just like pyramidal neurons of the olfactory cortex, KCs show sparse responses to odors^31,32^. The sparseness of the KC response depends at least in part on a single, key inhibitory neuron in the MB called anterior paired lateral (APL) neuron ^33^. Impairment of sparse coding by inactivating APL neuron causes learning defects, demonstrating that sparse coding at KCs is essential for stimulus-specific learning^33,34^. However, sparse coding alone is not enough to prevent unspecific association. First, there are significant overlaps between representations of odors whether or not they are chemically related^35^. The degree of such overlap directly correlates with the degree of crosstalk in plasticity^5^. Second, while, on average, only about 5 % of KCs reliably respond to multiple presentations of a given odor, there are up to 15 % additional KCs that unreliably respond in a given trial^32^. Those unreliable responders show smaller Ca^2+^ responses than reliable ones^32^. While the former phenomena (i.e. reliable overlap) could be beneficial for animals by contributing to biologically important generalization across stimuli^5,35^, the latter (i.e. unreliable overlap) will be only detrimental for learning by causing unspecific learning. This is important because dopamine-induced plasticity at KC-MBON synapses is so effective that even a single, brief odor-DA pairing can induce a long-lasting change in odor responses in some MBONs^5,28,29^. Furthermore, despite the prevailing model of coincidence detection of Ca^2+^ and DA signals by Rutabaga, DA alone can elevate cAMP level in KCs to some extent even without activation of KCs^27,36^. These considerations prompted us to search for additional mechanisms at synaptic terminals to prevent unspecific association.

Recently reported ultrastructural connectomics of the MB circuit revealed that there are surprisingly large numbers of KC-KC connections. In fact, more than 80% of local synaptic inputs to the KC axons are provided by the cholinergic KCs themselves^20^. Since no excitatory role was found for acetylcholine (ACh) in KC axons^37^, it suggests that ACh most likely acts as a neuromodulator rather than fast excitatory neurotransmitter. Although ACh is the main excitatory neurotransmitter in the *Drosophila* brain, its modulatory action has been understudied. The *Drosophila* brain expresses only two types of metabotropic muscarinic ACh receptors: mAChR-A, coupled to Gq, and mAChR-B, coupled to G_i/o_^38,39^. Because of the distinct downstream pathways, genetic manipulations of one of the receptors do not result in functional compensation by the other. Furthermore, mAChR-A is expressed only in KC dendrites^40^, suggesting a possible role for mAChR-B in KC-KC axonal neuromodulation.

Here we show that KC-KC axonal interaction is mediated by mAChR-B. This mAChR-B-mediated neuromodulation has two roles: it decreases both odor-evoked Ca^2+^ elevation and DA-induced cAMP elevation. Thus, this neuromodulation acts to reduce both signals that are required for KC-MBON synaptic plasticity. We further show that knocking down mAChR-B results in unspecific learning. That is, an odor that was not coupled to a punishment signal is perceived as being coupled to one. We therefore suggest that active KCs release ACh on cognate KCs that are less or non-active. This lateral neuromodulation enhances stimulus specificity of learning.

## Results

### γ KCs have extensive lateral axo-axonal connections

Our central hypothesis is that abundant KC-KC axonal interactions mediate essential neuromodulation for learning. We therefore used the recently published *Drosophila* brain connectome^41^ to examine KC-KC interactions. Our goal was to uncover which type of KCs has the most lateral connections and to examine the cellular compartment in which these lateral KC-KC connections occur. As expected, almost no KC-KC interactions are observed between KC dendrites, and the vast majority of KC-KC interactions are between KC axons (Figure S1A). There are three types of KCs (α/β, α’/β’ and γ), which are defined according to their innervation pattern^17,18^. αβ and α’β’ KCs bifurcate and send axons to the vertical (α and α’) and horizontal (β and β’) lobes, whereas γ KCs send axons only to the horizontal γ lobe. These different KCs play different roles in olfactory learning^26,42^. KC-KC interactions were mostly limited to within-subtypes (Figure S1A). γ KCs showed the most robust connectivity, with each KC synapsing on average with 190 other KCs, whereas α/β and α’/β’ synapsed on average on 145 and 109 KCs respectively (Figure S1B).

### mAChR-B is expressed in γ KC axonal terminals

Recently, KCs were found to be cholinergic^37^. It has been also suggested that KC axons are essentially devoid of nicotinic receptors, as local application of ACh or nicotine on KC axons generated no Ca^2+^ signal^37^. Although we recently showed that mAChR-A is expressed in KCs, it is localized at the dendrites^40^. As the fly brain only expresses two muscarinic receptors^43,44^, mAChR-B is the most logical candidate to mediate KC-KC axonal interactions. We therefore first examined which types of KCs express mAChR-B. To this end, we used two driver lines that can express GAL4 wherever mAChR-B is endogenously expressed; one is from the Minos-mediated integration cassette (MiMIC) collection^45^, and the other from the collection of T2A-GAL4 knockins^46^. The MiMIC insertion resides in the 5’-untranslated region of the mAChR-B gene, and we used recombinase-mediated cassette exchange (RMCE) to replace the original MiMIC cassette that contains GFP with a MiMIC cassette containing GAL4^45^ to generate a new mAChR-B-MiMIC-GAL4 driver line. In the case of the mAChR-B-T2A-GAL4 knockins, the T2A-GAL4 cassette is inserted immediately upstream of the mAChR-B stop codon. Enhancer-driven eGFP expression using those two independent lines was evident in α/β and γ but not α’/β’ KCs (Figure 1). Consistent with these results, single-cell or cell-type-specific transcriptomic analyses of the *Drosophila* brain^43,44,47^ suggest the mAChR-B is expressed more strongly in αβ and γ than in α’β’ KCs (Figure S2). To determine which subcellular compartments express mAChR-B, we generated a transgene of UAS-mAChR-B tagged with the hemagglutinin (HA) tag in its C-terminal and used the pan-KC driver line, OK107-GAL4. Overexpression of mAChR-B-HA revealed expression in the axonal lobes and cell bodies but not in the dendrites at the calyx (Figure 1). These results suggest that in the MB, mAChR-B is preferentially expressed in the axons of α/β and γ KCs.

**Figure 1:**
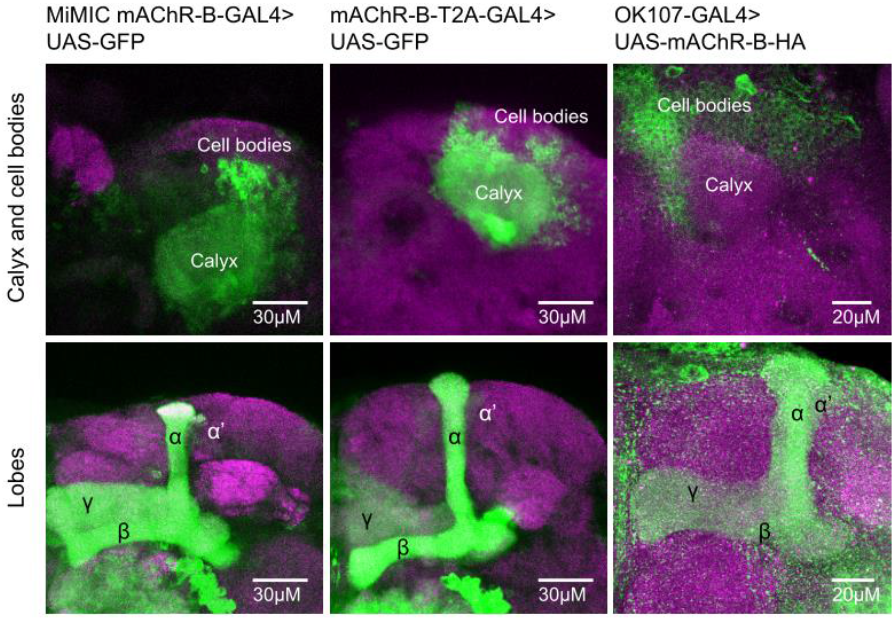
mAChR-B expression pattern. Maximum intensity projection of confocal sections through the central brain of a fly carrying MiMIC-mAChR-B-GAL4 (*left*) or mAChR-B-T2A-GAL4 (*middle*), and UAS-GFP transgenes. MB αβ and γ lobes are clearly observed (*bottom*, 80 and 71 confocal sections respectively, 1 μm). Very weak GFP expression is observed in α’β’ lobes. As expected from soluble GFP labeling the calyx is clearly observed (*top*, 44 and 24 confocal sections respectively, 1 μm). *Right*, the pan KC driver, OK107-GAL4 was used to overexpress UAS-mAChR-B-HA (with an HA tag). While KC axons at the MB lobes are clearly visible (*bottom*, 43 confocal sections, 0.5 μm), there is no expression at the calyx (compare to left and middle panels, 44 confocal sections, 0.5 μm) indicating mAChR-B is normally expressed in the axonal compartment.

### mAChR-B expression in KCs is required for efficient aversive olfactory learning

To test our central hypothesis, we next examined whether mAChR-B is required for aversive associative olfactory conditioning. To this end, we knocked down mAChR-B expression using one of two UAS-RNAi lines, “RNAi 1” or “RNAi 2,” the latter of which required co-expression of Dicer-2 (Dcr-2) for optimal knockdown (KD). To evaluate the RNAi efficiency, we performed quantitative real-time polymerase chain reaction (qRT-PCR). We knocked down mAChR-B expression using the pan-neuronal driver elav-GAL4. Both RNAi 1 and RNAi 2 significantly reduced mAChR-B levels, to about 55% and 25% of the original mAChR-B level (Figure S3). To knock down mAChR-B in KCs, we used the pan-KC driver OK107-GAL4. Short-term aversive memory was examined using two odors, 4-methylcyclohexanol (MCH) and 3-octanol (OCT), which are standard in the field^48^. For both UAS-RNAi transgenes, similar reduction in memory performance was observed, whether training was against MCH or OCT (Figure 2A). To verify that the reduced learning does not arise from defects in odor or electric shock perception, we examined naïve odor valence and preference as well as reaction to electric shock. Both UAS-RNAi transgenes had no effect on these properties (Figure S4). To rule out the possibility that developmental effects may underlie the learning defect observed in mAChR-B KD flies, we used tub-GAL80^ts^ to suppress RNAi 1 expression during development. GAL80^ts^ blocks the GAL4 transcription factor at 23°C but allows for normal expression of the UAS transgene at 31°C^49^. Flies were grown at 23°C until post-eclosion to block RNAi expression and allow for normal development and were then transferred to 31°C for 7 days to allow for the RNAi to be expressed and take effect. KD of mAChR-B only at the adult stage recapitulated the effect of constitutive KD of mAChR-B (Figure 2B). As a control to verify that GAL80^ts^ efficiently blocks RNAi expression, flies were constantly grown at 23°C (i.e., without transferring them to 31°C), thus blocking the RNAi expression also in adult flies. These flies showed normal olfactory associative learning (Figure 2B). Together, these results indicate that mAChR-B has a physiological role in associative olfactory aversive conditioning.

**Figure 2:**
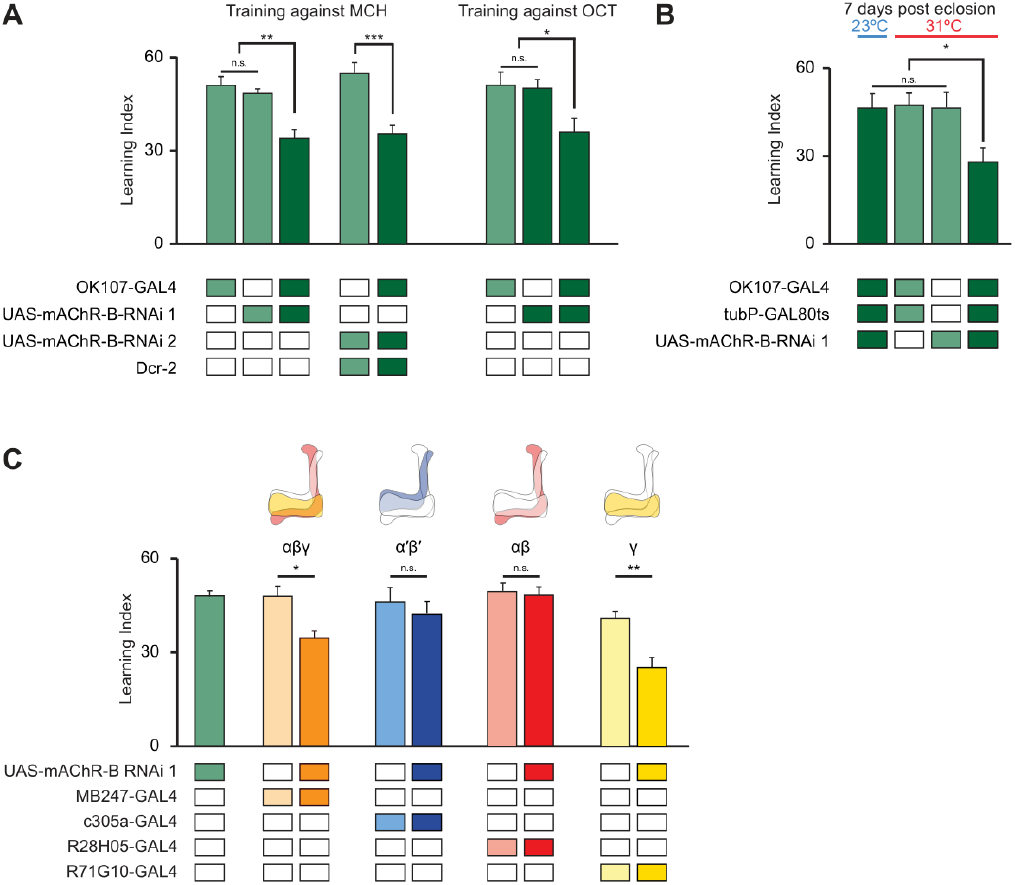
mAChR-B is required in γ KCs for short term aversive olfactory learning. **A.** Learning scores in flies with mAChR-B RNAi 1 or 2 driven by OK107-GAL4. mAChR-B KD reduced learning scores compared to controls (mean ± SEM), n (left to right): 49, 162, 77, 66, 61, 60, 63, 51; * p < 0.05, ** p<0.01, *** p<0.001; (Kruskal-Wallis with Dunn’s correction for multiple comparisons). For detailed statistical analysis see **Table S1**. **B.** Learning scores in flies with mAChR-B RNAi 1 driven by OK107-GAL4 with GAL80^ts^ repression. Flies raised at 23 °C and heated to 31 °C as adults (red horizontal bar) had impaired learning compared to controls. Control flies kept at 23 °C throughout (blue horizontal bar), thus blocking mAChR-B RNAi expression, showed no learning defects (mean ± SEM), n (left to right): 63, 60, 69, 61; * p < 0.05, ** p<0.01, *** p<0.001; (Kruskal-Wallis with Dunn’s correction for multiple comparisons). For detailed statistical analysis see **Table S1**. **C.** mAChR-B RNAi 1 was targeted to different subpopulations of KCs using c305a-GAL4 (α’β’ KCs), MB247-GAL4 (αβγ KCs), R28H05-GAL4 (αβ KCs), and R71G10-GAL4 (γ KCs). Learning scores mAChR-B RNAi 1 was expressed in αβ and γ KCs or γ KCs alone, but not when mAChR-B RNAi 1 was expressed in αβ or α’β’ KCs (mean ± SEM), n (left to right): 162, 73, 70, 54, 61, 72, 60, 91, 69; * p < 0.05, ** p<0.01, (Kruskal-Wallis with Dunn’s correction for multiple comparisons). For detailed statistical analysis see **Table S1**. The data for the UAS-mAChR-B RNAi 1 control are duplicated from panel A.

### mAChR-B is required for olfactory learning in γ KCs, but not in αβ or α’β’ KCs

Following the expression pattern of mAChR-B that is limited to αβ and γ KCs (Figure 1), we sought to examine in which KCs mAChR-B has a role in olfactory learning. To this end, we knocked down mAChR-B expression using RNAi 1 in different KC subtypes. As expected from the anatomical expression pattern of mAChR-B (Figure 1), aversive olfactory conditioning was impaired when mAChR-B was knocked down in αβ and γ KCs using MB247-GAL4, but not by KD in α’β’ KCs using c305a-GAL4 (Figure 2C). To examine if mAChR-B is required for aversive olfactory conditioning of both αβ and γ KCs, we used R28H05-GAL4 and R71G10-GAL4 lines, which drives expression only in αβ KCs or in γ KCs, respectively. Aversive olfactory conditioning was impaired when mAChR-B was knocked down in γ, but not αβ, KCs (Figure 2C). The combined expression pattern, anatomical connectivity, and behavioral results all point to a role of mAChR-B in the context of aversive olfactory conditioning, specifically in γ KCs axons.

### mAChR-B suppresses odor responses only in γ KC axons

To gain mechanistic understanding of the role played by mAChR-B in olfactory learning, we next examined whether mAChR-B contributes to olfactory responses in KCs. To this end, we expressed the genetically encoded Ca^2+^ indicator, GCaMP6f^50^, with or without mAChR-B RNAi 1 using MB247-GAL4 (labels α/β and γ KCs), c305a-GAL4 (labels α’/β’ KCs), or R71G10-GAL4 (labels only γ KCs) and examined using *in vivo* 2-photon functional imaging the effect mAChR-B KD has on responses to MCH and OCT. To check for various effects on the odor response, we examined three different parameters: the peak of the “On” and “Off” responses and the overall strength of the odor response as measured by the integral of the response (see methods). Following mAChR-B KD, we found a significant and reliable increase in odor responses in γ KCs (Figure 3). The increase was observed in all three parameters. Importantly, consistent with our observation that mAChR-B is expressed in the axons, no effect was observed in the dendritic arborizations in the calyx. This was true for both the broad MB247-GAL4 driver line and the γ-KC-specific driver line, R71G10-GAL4 (Figure 3). The fact that no effect was observed in KC dendritic arborization (Figure 3) indicates that KC recruitment by upstream projection neurons is not impaired by mAChR-B KD and that mAChR-B neuromodulation occurs locally at KC presynaptic axonal terminals. To test whether the observed increase in Ca^2+^ response affects synaptic release, we used the genetically encoded ACh sensor, ACh3.0^51^ that reports ACh level in the synaptic cleft. We expressed ACh3.0 in KCs using the MB247-GAL4 driver line and examined KC synaptic release following odor application (MCH and OCT). Consistent with the above result, a significant increase of all three examined parameters (as above) was observed also for released ACh levels (Figure S5).

**Figure 3:**
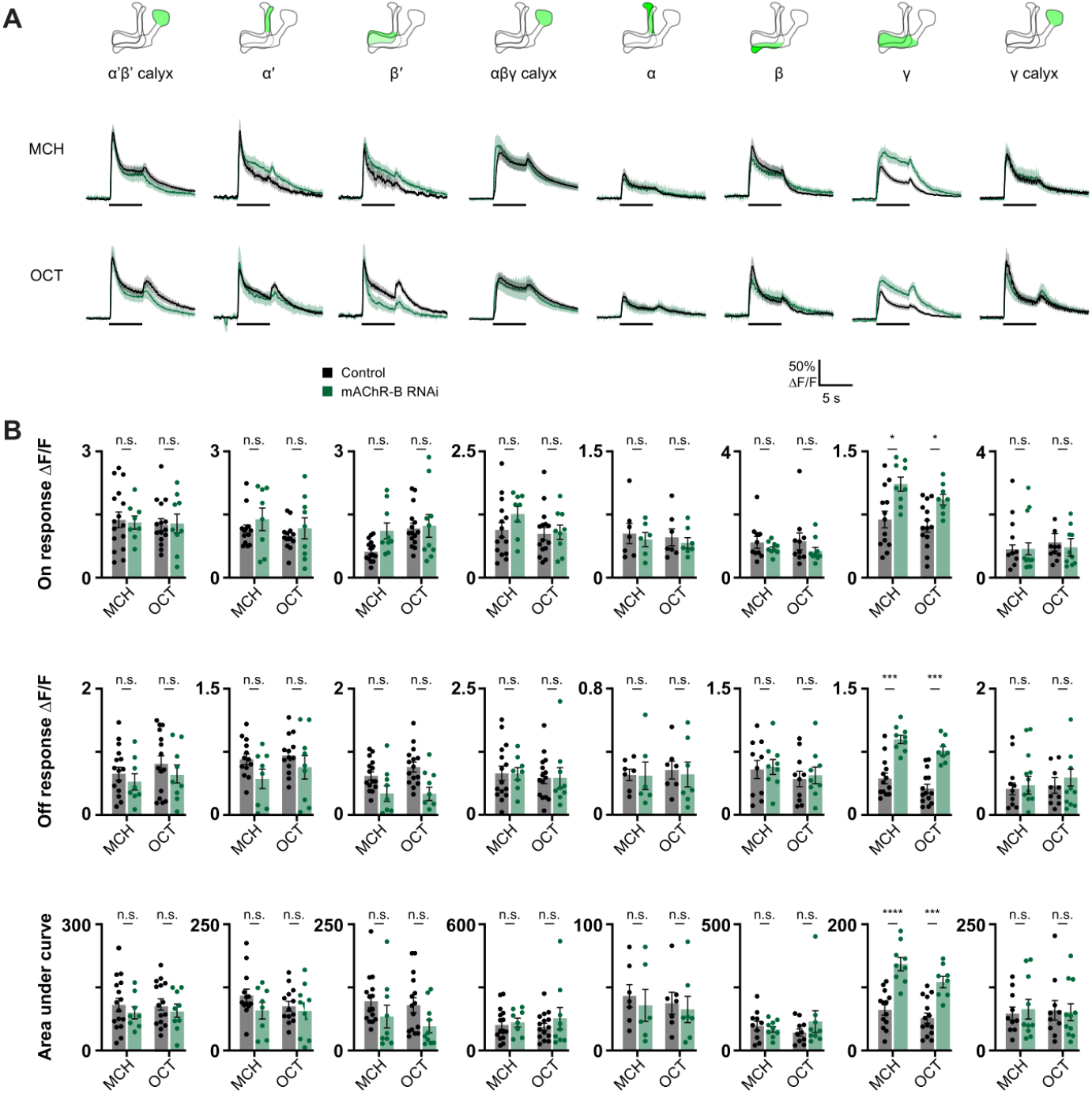
mAChR-B knockdown increases odor responses only in γ KC axons. Odor responses to MCH and OCT were measured in control flies (GAL4>UAS-GCaMP6f) and knockdown flies (GAL4>UAS-GCaMP6f, UAS-mAChR-B RNAi 1). The following driver lines were used c305a-GAL4 (α’β’ KCs), MB247-GAL4 (αβγ KCs), and R71G10-GAL4 (labels only γ KCs). **A.** ΔF/F of GCaMP6f signal in different areas of the MB in control (black) and knockdown (green) flies, during presentation of odor pulses (horizontal lines). Data are mean (solid line) ± SEM (shaded area). Diagrams illustrate which region of the MB was analyzed. **B.** Peak “on” response (top), Peak “off” response (middle), and the integral of the odor response (bottom) of the traces presented in A (mean ± SEM). Only in γ KC axons a significant increase in odor responses is observed. This increase was observed for both odors and in all modes of analysis. n for control MCH, OCT and KD MCH, OCT flies, respectively: α’β’ calyx, 16, 15, 8, 9; α’, 13, 13, 8, 9; β’, 15, 15, 9, 10; αβγ calyx, 15, 15, 8, 9; α, 7, 7, 6, 7; β, 10, 10, 9, 9; γ, 13, 14, 9, 8; γ calyx, 10, 10, 10, 11. * p < 0.05, *** p < 0.001, **** p < 0.0001 (Mann-Whitney test with Holm Šídák correction for multiple comparisons). For detailed statistical analysis see **Table S1**.

We next examined whether overexpression of mAChR-B has an opposite effect to that of mAChR-B KD. To this end, we generated a UAS-mAChR-B transgene and used R71G10-GAL4 to drive mAChR-B in γ KCs. To verify the efficiency of the UAS-mAChR-B overexpression, we used the pan-neuronal driver elav-GAL4 to overexpress mAChR-B and examined its level using qRT-PCR. Overexpression of mAChR-B resulted in increased levels of about 155% relative to the original level (Figure S3). As expected from the above results (Figs. 1 and 3), overexpression of mAChR-B resulted in decreased odor response in γ KCs axons but not in the dendritic arborization (Figure 4). Taken together, these results suggest that mAChR-B normally acts to reduce the odor-evoked Ca^2+^ response and synaptic output at γ KC axons.

**Figure 4:**
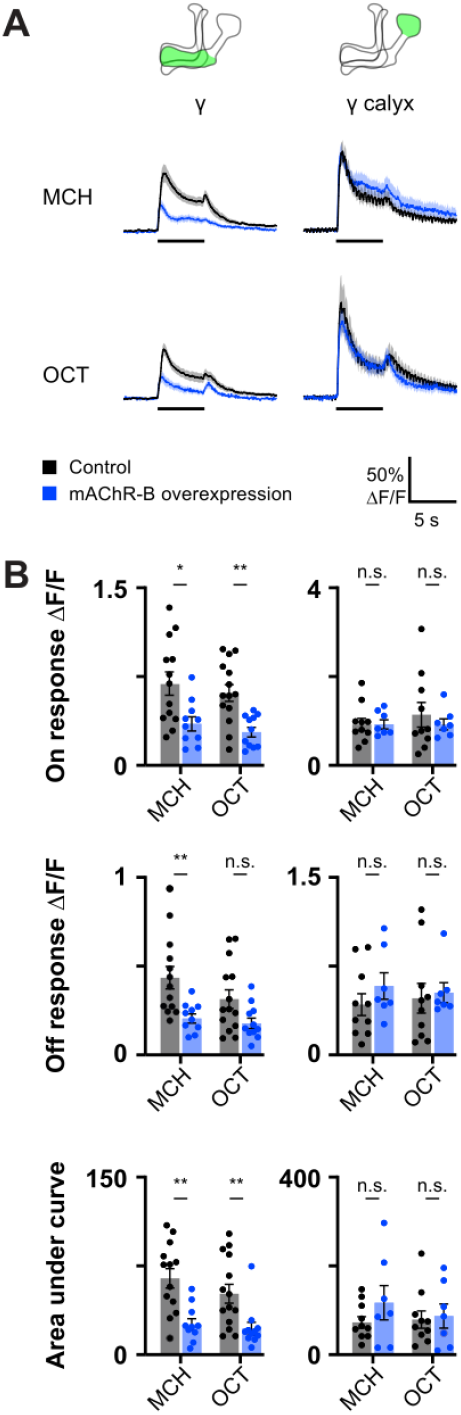
mAChR-B overexpression decreases odor responses only in γ KC axons. Odor responses to MCH and OCT were measured in control flies (R71G10-GAL4>UAS-GCaMP6f) and overexpression flies (R71G10-GAL4>UAS-GCaMP6f, UAS-mAChR-B). **A.** ΔF/F of GCaMP6f signal in the calyx and lobe of γ KCs for control (black) and overexpression (blue) flies, during presentation of odor pulses (horizontal lines). Data are mean (solid line) ± SEM (shaded area). Diagrams illustrate which region of the MB was analyzed. **B.** Peak “on” response (top), Peak “off” response (middle), and the integral of the odor response (bottom) of the traces presented in A (mean ± SEM). Only in γ KC axons a significant decrease in odor responses is observed. This decrease was observed for both odors and in all modes of analysis. n for control MCH, OCT and overexpression MCH, OCT flies, respectively: γ, 14, 13, 10, 11; γ calyx, 10, 10, 7, 7. * p < 0.05, ** p < 0.01, (Mann-Whitney test with Holm Šídák correction for multiple comparisons). For detailed statistical analysis see **Table S1**.

### mAChR-B suppresses dopamine-induced increase in cAMP

Studies in heterologous expression systems suggested that mAChR-B is coupled to G_i/o_^38,39^. So we next examined whether ACh application on KCs affects cAMP level. To this end, we used MB247-GAL4 to drive expression of the recently developed genetically encoded cAMP indicator, cAMPr^52^. We used tetrodotoxin (TTX) to block circuit effects and locally applied ACh (1 mM) using a puff pipette targeted to KC axons at the horizontal lobe. We observed that ACh application significantly reduces cAMP level (Figure 5A, black). To verify that this reduction indeed arises from activation of mAChR-B expressed in KCs, we repeated the experiment but with mAChR-B KD in KCs using RNAi 1. KD of mAChR-B almost completely suppressed the reduction in cAMP level (Figure 5A, green). Thus, even if there were residual circuit effects in the presence of TTX (e.g. via the APL neuron), KD of mAChR-B in KCs should not have affected these residual effects. Together, the effect of mAChR-B KD in KCs clearly indicates that the net effect of ACh on cAMP signaling in KCs is inhibitory and that it is mediated by mAChR-B.

**Figure 5:**
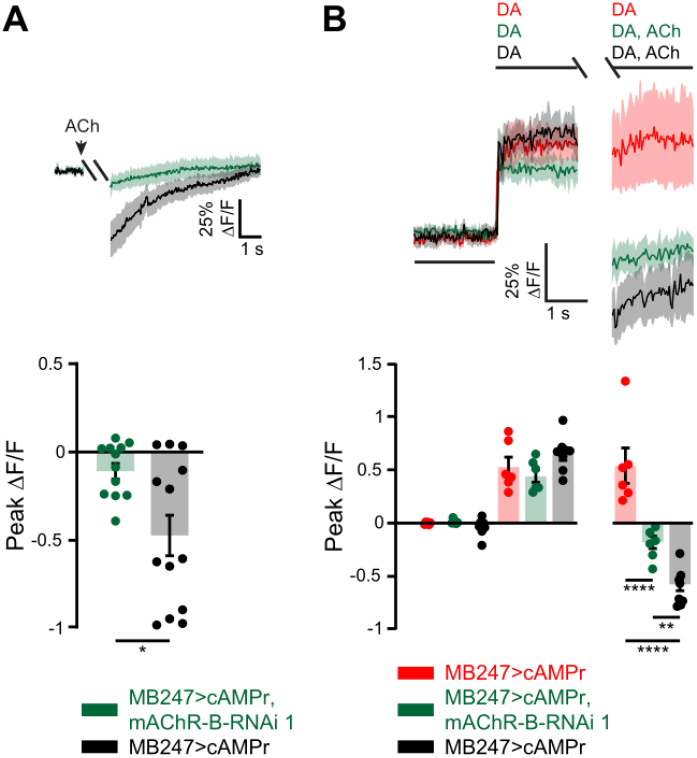
mAChR-B decreases cAMP level. cAMP level was measured using the single-wavelength fluorescent sensor for cyclic AMP, UAS-cAMPr. MB247-GAL was used to drive expression. **A.** *Top*, ΔF/F of cAMPr following activation of mAChR-B using a 2 s puff (gap) of 1mM ACh (black). To abolish any circuit effects 1 μM TTX was bath applied. KD of mAChR-B using UAS-mAChR-B RNAi 1 (green) abolished the decrease in cAMP, indicating that the observed cAMP decrease indeed arises from mAChR-B activation. Data are mean (solid line) ± SEM (shaded area). *Bottom*, Peak response of the top presented traces (mean ± SEM). n for *wt* and KD flies respectively, 13, 12; * p < 0.05, (Mann-Whitney two-tailed rank test). For detailed statistical analysis see **Table S1**. **B.** *Top*, ΔF/F of cAMPr. Application of 5 mM DA resulted in sustained increase in cAMP levels (red). The 2 s gap is when a puff of either DA or ACh and DA together was given. Activation of mAChR-B using a puff of 1mM ACh (black) significantly decreased cAMP levels even below the initial level. To abolish any circuit effects 1 μM TTX was bath applied. KD of mAChR-B using UAS-mAChR-B RNAi 1 (green) resulted in a less severe decrease of cAMP compared to the *wt* conditions (black), indicating that the observed cAMP decrease indeed arises from mAChR-B activation. The residual decrease in cAMP level most probably arises from the incomplete KD of mAChR-B using mAChR-B RNAi 1 (Figure S3). Data are mean (solid line) ± SEM (shaded area). *Bottom*, Peak response of the top presented traces (mean ± SEM). n for DA alone, DA with mAChR-B activation, and for KD flies respectively, 6, 8, 6; * p < 0.05, (Shapiro-Wilk normality test, followed by two-way ANOVA with Tukey’s correction for multiple comparisons). For detailed statistical analysis see **Table S1**.

DA exerts its main effect on aversive olfactory conditioning by activation of Dop1R1, which is Gs coupled, and increases cAMP level^27,36^. We therefore examined whether activation of mAChR-B can counter the effect of DA application on cAMP level. As before, MB247-GAL4 was used to drive cAMPr and TTX was used to block any circuit effects. DA at high concentration (5 mM) was bath applied and generated a robust and sustained elevation in cAMP level (Figure 5B, red). Local application of ACh significantly reduced cAMP to below the initial basal level (Figure 5B, black). To verify that this reduced level of cAMP indeed arises from mAChR-B activation, we knocked down mAChR-B in KCs using RNAi 1. Under these conditions, ACh application reduced cAMP level but only to the initial basal level (Figure 5B, green). Recalling that RNAi 1 reduced mAChR-A to only about 55% of initial level, it is more than likely that the remaining mAChR-B underlie the decrease in cAMP observed. Nevertheless, mAChR-B KD attenuated ACh-induced decrease in cAMP (Figure 5B, Table S1). Together, these results indicate that mAChR-B can counter the DA-induced increase in cAMP by reducing cAMP level.

### mAChR-B mediates lateral KC neuromodulation

The results thus far demonstrate that mAChR-B can counter both signals required for efficient plasticity in KC presynaptic axonal terminals: Ca^2+^ and cAMP elevation. We now turn to examine the source of ACh that activates KC mAChR-B. The vast KC-KC axonal interactions (Figure S1) suggest that KCs are the source of the ACh. We therefore used MB247-GAL4 to express the tetanus toxin light chain (TNT), which inhibits synaptic transmission^53^. Blocking KC synaptic release abolished the effect of mAChR-B KD on odor-evoked Ca^2+^ responses (Figure S6), suggesting that KCs are the source of the ACh that attenuate odor-evoked Ca^2+^ responses in normal flies.

The existence of lateral KC-KC axonal interactions and axonal localization of mAChR-B suggest that KCs affect other, neighboring KCs via mAChR-B. However, mammalian muscarinic M2 that is also coupled to G_i/o_ often acts as an autoreceptor^54,55^. Therefore, to examine whether mAChR-B acts as an autoreceptor or promotes lateral neuromodulation, we sought to activate a subpopulation of KCs and examine how synaptic release from this subpopulation affects odor responses in other cognate KCs (Figure 6A). To this end, we used the recently developed Sparse Predictive Activity through Recombinase Competition (SPARC) method ^56^ to express the red-light-activated channel, CsChrimson^57^ in a subpopulation of KCs, thus allowing for their optogenetic activation (Figure 6A). By expressing nSyb-PhiC31 recombinase pan-neuronally together with intermediate SPARC2-CsChrimson::tdTomato variants (i-intermediate) and MB247-GAL4, we generated flies that expressed CsChrimson::tdTomato in approximately 15% of α/β and γ KCs in a stochastically distributed manner. GCaMP6f was expressed in all α/β and γ KCs using MB247-LexA and LexAop-GCaMP6f (Figure 6B). This experimental configuration allows us to examine how activation of a subpopulation of KCs affects odor responses of other KCs (Figure 6A). To ensure that only lateral effects are measured, we examined odor responses only in KCs that did not express CsChrimson (Figure S7). KC odor responses were examined with and without optogenetic activation (Figure 6A). Consistent with the inhibitory effect mAChR-B had on KC odor responses (Figs. 3 and 4), optogenetic activation of the CsChrimson resulted in decreased odor responses in CsChrimson-negative KCs, demonstrating a lateral suppression of KC activity (Figures 6C-6H). Furthermore, consistent with the above results (Figs. 3 and 4), this lateral suppression was completely abolished with mAChR-B KD in all α/β and γ KCs (Figures 6I-6N). Thus, these results demonstrate that mAChR-B mediates lateral KC-KC neuromodulation.

**Figure 6:**
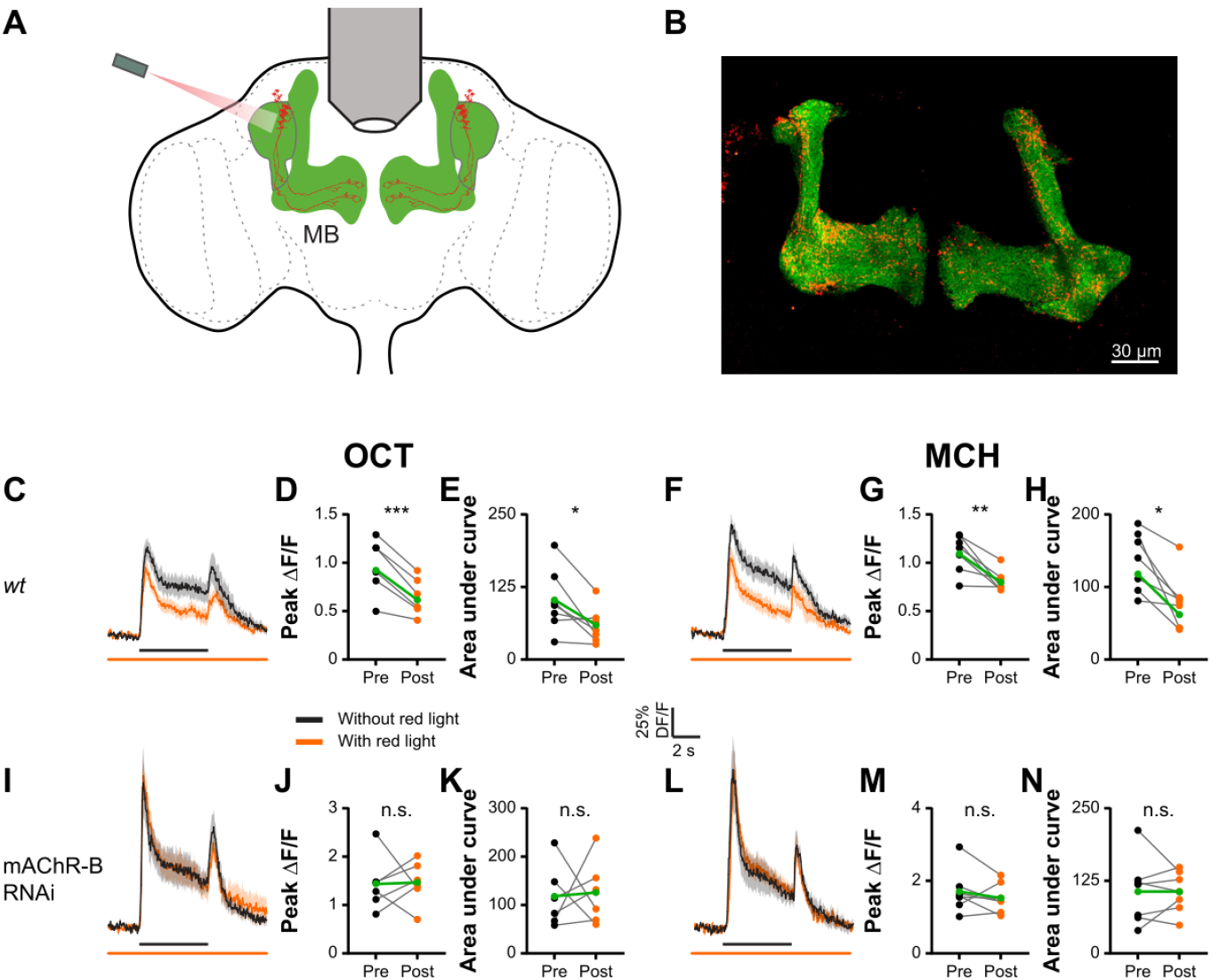
Axo-axonal lateral neuromodulation underlies mAChR-B effects. **A.** Experimental configuration. GCaMP6f was expressed in all αβγ KCs. The SPARC method was used to drive CsChrimson in ~15% of αβγ KCs. Responses to MCH and OCT of γ KCs that do not express CsChrimson were examined with and without optogenetic activation of the ~15% KC subpopulation. **B.** Example of a fly brain with the strategy presented in A. Maximum intensity projection of 28 confocal sections (1 μm) through the central brain of a fly carrying MB247-LexA-LexAop-GCaMP6f, and CsChrimson::tdTomato in stochastically distributed subsets of neurons within the MB247-GAL4 driver line transgenes. **C, F, I, L.** ΔF/F of GCaMP6f signal in the γ KC lobe for control (C, F) and KD (I, L) flies, during presentation of odor pulses (OCT, C, I; MCH, F, L; horizontal black lines) without (black) or with (orange) optogenetic activation of the ~15% KC subpopulation expressing CsChrimson. MB247-LexA was used to drive LexAop-GCaMP. MB247-GAL4 was used to drive UAS-mAChR-B RNAi 1 and TI{20XUAS-SPARC2-I-Syn21-CsChrimson::tdTomato-3.1}CR-P40. Data are mean (solid line) ± SEM (shaded area). GCaMP signals were taken from regions not expressing CsChrimson (see Figure S6). **D, E, G, H, J, K, M, N.** Peak “on” response (D, G, J, M), and the integral of the odor response (E, H, K, N) of the traces presented in C, F, I, L before (left) or after (right) optogenetic activation. A significant decrease in odor responses is observed only in control flies indicating lateral neuromodulation. This lateral neuromodulation is absent in KD flies not expressing mAChR-B. n for control and KD flies, respectively: OCT, 7, 6; MCH, 7, 7. * p < 0.05, ** p < 0.01, *** p < 0.001, (Wilcoxon matched-pairs signed rank test). For detailed statistical analysis see **Table S1**.

### mAChR-B decreases unspecific learning

How can the physiological actions of mAChR-B we demonstrated so far explain the impaired learning in mAChR-B KD flies (Figure 2)? In the MB, coincidence of KC activity (i.e. Ca^2+^ increase), which represents olfactory signal, and DA input (i.e. cAMP increase), which represents reinforcement signal, is considered to induce synaptic plasticity and learning. If mAChR-B in KCs acts synergistically with dopamine signaling pathway, that could explain the impaired learning in mAChR-B KD flies (i.e. defective learning). However, our results show antagonistic actions of ACh and DA in KCs (Figure 5), which makes this scenario unlikely. As an alternative scenario, we hypothesized that lateral KC-KC interactions mediated by mAChR-B may prevent unspecific learning by suppressing both Ca^2+^ and cAMP increase in less active or inactive KCs during conditioning. These two scenarios predict entirely different types of learning impairment (i.e. defective versus unspecific learning; Figure S8) in mAChR-B KD flies with respect to the valence change in paired odor (conditioned stimulus, CS^+^) and unpaired, control odor (CS^-^). That is, in defective learning, the negative shift of the valence of CS^+^ after aversive learning should be diminished (Figure S8C), whereas in unspecific learning, the valence shift of CS^+^ should not be affected, but CS^-^ should show more negative shift than normal (Figure S8D).

To discriminate these possibilities, we trained flies with a 1-minute training protocol, as was done for the above behavior experiments (see methods) and then examined the changes in odor valences. For this set of experiments, flies were conditioned against MCH, and the valence of either MCH or OCT was examined (Figure 7A). As expected, following aversive conditioning against MCH, MCH became more aversive than it was before (Figure 7B). For the parental control flies, following aversive conditioning against MCH, OCT became slightly more attractive (Figure 7B). In agreement with our findings that mAChR-B opposes DA neuromodulation and that KCs exert lateral neuromodulation on cognate KCs, KD of mAChR-B had no effect on the increase in aversion towards MCH (Figure 7B). However, the change in OCT valence reversed, and rather than becoming more attractive, OCT became more aversive (Figure 7B). These results indicate that in the absence of mAChR-B and lateral KC-KC neuromodulation, OCT undergoes unspecific conditioning (Figure 7B), consistent with the unspecific learning model suggested above (Figure S8).

**Figure 7:**
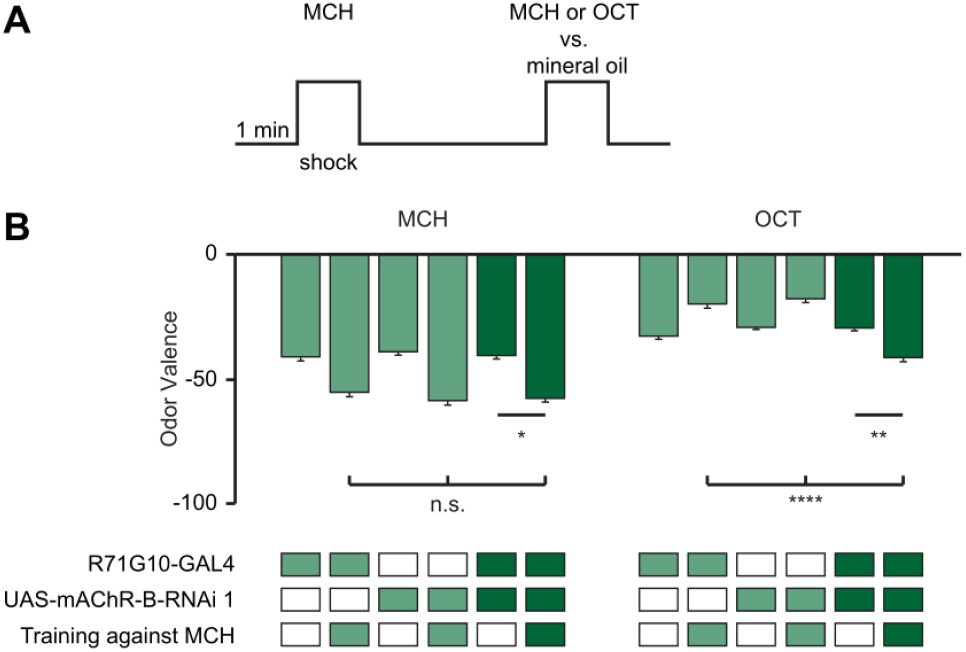
mAChR-B KD results in unspecific conditioning. **A.** Experimental protocol. Flies were conditioned against MCH using 12 equally spaced 1.25 s electric shocks at 50 V. Flies were then subjected to OCT or MCH for valence evaluation (see methods). **B.** MCH or OCT valence (as designated) observed with or without pre-exposure to conditioning against MCH in flies with mAChR-B RNAi 1 driven by R71G10-GAL4 (γ KCs). Following conditioning against MCH both mAChR-B KD and control flies showed the same increase in aversion towards MCH. In contrast, when OCT valence was examined, mAChR-B KD flies showed increased aversion towards OCT whereas the parental controls showed reduced aversion towards OCT (mean ± SEM), n (left to right): MCH: 63, 61, 65, 70, 73, 75; OCT: 96, 76, 66, 77, 101, 76; * p < 0.05, ** p<0.01, *** p<0.001; (Kruskal-Wallis Dunn’s correction for multiple comparisons). For detailed statistical analysis see **Table S1**.

## Discussion

Here we showed that KC-KC axonal interaction is mediated by mAChR-B. This mAChR-B-mediated neuromodulation has dual roles: it decreases both odor-evoked Ca^2+^ elevation and DA-induced cAMP elevation. Thus, this neuromodulation suppresses both signals that are required for KC-MBON synaptic plasticity. In behavior experiments, we demonstrated that mAChR-B KD in KCs impairs stimulus specificity of learning. Our study reveals a novel form of local neuromodulation, which improves sensory discrimination during learning.

In this study, we identified the first biological functions, to our knowledge, of axo-axonic synapses between KCs. Olfactory coding in the insect MB is a well-established model system to study the circuit mechanisms and benefits of sparse sensory representations. The abundance of KC-KC synapses at the axons discovered by the EM connectome^20^ surprised the field at first because excitatory cholinergic interactions may ruin the very benefit of the sparse coding in olfactory learning. However, our Ca^2+^ imaging demonstrated that the net effect of those cholinergic transmissions is, in fact, inhibitory. The lateral inhibition mediated by mAChR-B should further enhance, rather than ruin, the benefit of sparse coding and thereby improve the stimulus specificity of learning. Even though the population of KCs that show reliable responses to a given odor is sparse (~5 %), many more KCs are activated in a given odor presentation. This is because there is a larger population of unreliable responders, making up to ~15 % of total KCs active in a given trial^32^. Since those unreliable responders tend to show weaker Ca^2+^ responses than the reliable ones, it is reasonable to speculate that mAChR-B-mediated mutual inhibition would preferentially suppress unreliable responders, letting reliable responders win the lateral competition. Since even a single, 1-second odor-DAN activation pairing can induce robust KC-MBON synaptic plasticity^5^, presence of unreliable responders can significantly compromise the synapse specificity of plasticity. Restricting Ca^2+^ responses to reliable responders should therefore greatly enhance the stimulus specificity of learning.

To support our finding, selective inhibition of Go signaling in KCs by expressing pertussis toxin (PTX) impairs aversive learning, and this effect was mapped to α/β and γ KCs^58^, which we found to express mAChR-B most abundantly. Furthermore, expression of PTX disinhibits odor-evoked vesicular release in γ KCs, and PTX-induced learning defect was ameliorated by hyperpolarization or blocking synaptic output of γ KCs^59^. We argue that mAChR-B-mediated inhibitory communication between γ KCs contributes at least in part to those previous observations.

Lateral communication through mAChR-B also suppresses cAMP signals in KCs, which counteracts Dop1R1-mediated DA action during associative conditioning. Since DA release in the MB likely takes a form of volume transmission^20^, it cannot provide target specificity of modulation. Furthermore, while induction of LTD depends on coincident activity of KCs and DANs^5,28–30^, elevation of cAMP can be triggered by DA application alone^27^, although DA input followed by KC activity could induce opposite plasticity (i.e. potentiation) via another type of DA receptor^29^. Thus, lateral inhibition of cAMP signals by G_i/o_-coupled mAChR-B plays an essential role in the maintenance of target specificity of modulation. Taken together, dual actions of mAChR-B on local Ca^2+^ and cAMP signals at KC axons, where plasticity is supposed to take place, should directly contribute to synapse specificity of plasticity (Figure S9). If animals lack mAChR-B in KCs, axons of unreliable responders to CS^+^ would stay mildly active during conditioning. Furthermore, DA release on KCs causes some unchecked increase in cAMP in inactive and mildly active KCs. Consequently, some plasticity occurs in these KCs, even if to a lesser extent than in the KCs that are reliably and strongly activated by the CS^+^. Thus, absence of mAChR-B would minimally affect plasticity of KCs that are reliably activated by the CS^+^, assuming that those KCs are nearly maximally depressed in the presence of mAChR-B. However, other KCs, which may include reliable responders to the CS^-^, will also undergo plasticity (Figure S9). This should result in unspecific association, and that is exactly the type of learning defect we observed in mAChR-B KD flies (Figure 7).

What may be the cellular mechanisms underlying the effects of mAChR-B on cAMP and Ca^2+^ level? mAChR-B was shown to be coupled to G_i/o_^39^, which is known to inhibit the cAMP synthase, adenylate cyclase^60^, which is widely expressed in KCs^23,27^. In addition, the G_βγ_ subunits have been demonstrated to be able to directly block voltage-gated Ca^2+^ channels^61^. G_βγ_ can also directly open inward rectifying potassium channels^62^ that would oppose the changes in membrane potential required for the gating of voltage-gated Ca^2+^ channels, although these potassium channels are not broadly expressed in KCs^43,44,47^.

Although the majority of study on population-level sensory coding has focused on somatic Ca^2+^ or extracellular electrophysiological recordings, our study sheds light on the importance of local regulation of Ca^2+^ and other intracellular signals at the axons when it comes to stimulus specificity of learning. Are there other mechanisms that may be involved in reducing unspecific conditioning? One potential source of such mechanisms is the APL neuron, a single GABAergic neuron in the MB that is excited by KCs and provides feedback inhibition to KCs^33,63–65^. Since activity of APL neuron contributes to sparse and decorrelated olfactory representations in KCs^33^, it is possible that GABAergic input to KC axons also serves to prevent unspecific learning. Release of GABA onto KC axons is expected to have similar effects as the activation of mAChR-B. Specifically, the activation of the G_i/o_-coupled GABA-B receptors that are widely expressed in KCs^43,44,47^, should have similar effects as activation of mAChR-B. However, in our experiments, lateral inhibition induced by optogenetic activation of a subset of KCs was completely suppressed by mAChR-B KD (Figure 6), suggesting that APL neuron did not contribute to lateral suppression of Ca^2+^ response at least in our experimental condition. This result is consistent with the prediction that individual KCs inhibit themselves via APL neurons more strongly than they inhibit the others due to the localized nature of the activity of APL neuron’s neurites and the geometric arrangement of the ultrastructurally identified synapses^66^. Nonetheless, whether APL neuron contributes to sparsening of axonal activity to prevent unspecific conditioning remains to be examined. In summary, the current study identifies functional roles of axo-axonic cholinergic interactions by uncovering previously unknown local neuromodulation that can enhance the stimulus specificity of learning, and refines the dopamine-centric view of MB plasticity.

## Acknowledgments

We thank the Bloomington Drosophila Stock Center (NIH P40OD018537), the Vienna Drosophila RNAi Center, Dr. Andrew Lin (University of Sheffield), Dr. Justin Blau (New York University), and Dr. Oren Schuldiner (Weizmann Institute of Science) for providing Drosophila strains. We also thank the Drosophila Genomics Resource Center (supported by NIH grant 2P40OD010949) for cDNA clones. We thank Dr. Andrew Lin also for constructive comments on the manuscript. MP was supported by the Israel Science Foundation (ISF, 343/18) and the United States-Israel binational Science Foundation (BSF, 2019026, 2020636). TH was supported by NSF (2034783), BSF (2019026) and NIH (R01DC018874). AMD was supported by NIH (F32MH125582).

## Author contributions

JEM: Conceptualization, methodology, investigation, formal analysis, writing–review & editing, visualization. AMD: Methodology, investigation, writing–original draft, writing–review & editing. TH: methodology, investigation, formal analysis, writing–original draft, writing–review & editing, supervision, funding acquisition. MP: Initiated the project, conceptualization, methodology, investigation, formal analysis, software, writing–original draft, writing–review & editing, visualization, supervision, funding acquisition.

## STAR Methods

### RESOURCE AVAILABILITY

#### Lead Contact

Further information and requests for resources and reagents should be directed to and will be fulfilled by the lead contact, Moshe Parnas.

#### Materials Availability

Flies generated for this paper, data and code used to generate the figures will be available upon request. Requests should be directed to and will be fulfilled by the lead contact, Moshe Parnas.

#### Data and Code Availability

The data and code used to generate Figures 1-7, supplementary Figures S1-S9, and Tables S1, are available from the corresponding author upon request. The study did not generate any new code or dataset. Any additional information required to reanalyze the data reported in the paper is available from the lead contact upon request.

### EXPERIMENTAL MODEL AND SUBJECT DETAIL

#### Fly Strains

Fly strains were raised on cornmeal agar under a 12 h light/12 h dark cycle and studied 7–10 days post-eclosion. Strains were cultivated at 25 °C. In cases where a temperature-sensitive gene product (GAL80^ts^) was used, the experimental animals and all relevant controls were grown at 23 °C. To allow expression of RNAi with GAL80^ts^, experimental and control animals were incubated at 31 °C for 7 days. Subsequent behavioral experiments were performed at 25 °C.

Experimental animals carried transgenes over Canton-S chromosomes where possible to minimize genetic differences between strains. The following transgenes were used: UAS-GCaMP6f (BDSC #42747), MB247-lexA-lexAop-GCaMP6f (a gift from Dr. Andrew Lin), UAS-mAChR-B RNAi 1 (TRiP. HMS05691, Bloomington BDSC #67775), UAS-mAChR-B RNAi 2 (VDRC ID #107137), UAS-Dcr-2 (Bloomington BDSC #24651), tub-GAL80ts (BDSC #7108), MB247-GAL4 (BDSC #50742), OK107-GAL4 (BDSC #854), c305a-GAL4 (BDSC #30829), TI{2A-GAL4}mAChR-B[2A- GAL4] (BDSC #84650), MiMIC-mAChR-B-GAL4 (generated in-house, as previously described ^40^), elav-GAL4 (BDSC #458), UAS-TNT (BDSC #28838), nSyb-IVS-PhiC31 (BDSC #84151), TI{20XUAS-SPARC2-I-Syn21-CsChrimson::tdTomato-3.1}CR-P40 (BDSC #84144), GMR71G10-GAL4 (BDSC #39604), GMR28H05-GAL4 (BDSC #49472), UAS-GACh3.0 (BDSC #86549), UAS-cAMPr (a gift from Dr. Justin Blau), UAS-mCD8-GFP (BDSC #32186). UAS-mAChR-B and UAS-mAChR-B-HA were made in-house. Briefly, *Drosophila* mAChR-B was subcloned from a *Drosophila* Genomics Resource Center clone (DGRC, GEO08261) into the pBID-UASc plasmid using standard methods (Epoch Life Sciences Inc.). Transgenic strains were established by injecting pBID-UASc-mAChR-B constructs into attP40 landing site (BestGene. Inc.).

### METHOD DETAILS

#### Behavioral Analysis

For behavior experiments, custom-built, fully automated apparatus were used. Single flies were placed in clear polycarbonate chambers (length 50 mm, width 5 mm, height 1.3 mm) with printed circuit boards (PCBs) at both floors and ceilings. Solid-state relays (Panasonic AQV253) connected the PCBs to a 50 V source.

Mass flow controllers (CMOSens PerformanceLine, Sensirion) were used to control the airflow. An odor stream (0.3 l/min) was obtained by circulating the airflow through vials filled with a liquid odorant and was combined to a carrier flow (2.7 l/min) yielding a 10 fold dilution from the odor source. The odor source was prepared at 10 fold dilution in mineral oil. Together, a final 100 fold dilution of odors was used. Fresh odors were prepared daily. Two identical odor delivery systems delivered odors independently to each half of the chamber. The 3 l/min total flow, consisting of the carrier flow and the odor stimulus flow, was split between 20 chambers. Thus, each half chamber received a flow rate of 0.15 l/min per half chamber. The airflow from the two halves of the chamber converged at a central choice zone.

The 20 chambers were stacked in two columns of 10 chambers. The chambers were backlit by 940 nm LEDs (Vishay TSAL6400). Images were obtained by a MAKO CMOS camera (Allied Vision Technologies) equipped with a Computar M0814-MP2 lens. The apparatus was operated in a temperature-controlled incubator (Panasonic MIR-154) maintained at 25 °C.

A virtual instrument written in LabVIEW 7.1 (National Instruments) extracted fly position data from video images and controlled the delivery of odors and electric shocks. Data were analyzed in MATLAB 2015b (The MathWorks) and Prism 6 (GraphPad).

A fly’s odor preference was calculated as the percentage of time that it spent on one side of the chamber. Three behavior protocols were used in this study as previously described ^40,67,68^. (i.) For odor valence, an odor was delivered to one side of the chamber and mineral oil to the other side for two minutes, and then the sides were switched. (ii.) For conditioning experiments, OCT and MCH were presented for two minutes from each side of the chamber. This was followed by a single training session of one minute (see below). Following the training session, flies were allowed to recover for 15 minutes and then they were tested again for preference between OCT and MCH. (iii.) For nonspecific learning, flies were conditioned against MCH for one minute. This was followed by three minutes of recovery, and the valence of either OCT or MCH was evaluated as in point (i.) The naïve avoidance index was calculated as (preference for left side when it contains air) – (preference for left side when it contains odor). During training, MCH or OCT were paired with 12 equally spaced 1.25 s electric shocks at 50 V (CS^+^). The learning index was calculated as (preference for CS^+^ before training) – (preference for CS^+^ after training). Flies were excluded from analysis if they entered the choice zone fewer than 4 times during odor presentation.

#### Functional Imaging

Brains were imaged by two-photon laser-scanning microscopy (DF-Scope installed on an Olympus BX51WI microscope, Sutter). Flies were anesthetized on ice then a single fly was moved to a custom built chamber and fixed to aluminum foil using wax. The cuticle and trachea were removed in a window overlying the required area. The exposed brain was perfused with carbogenated solution (95% O_2_, 5% CO_2_) containing 103 mM NaCl, 3 mM KCl, 5 mM trehalose, 10 mM glucose, 26 mM NaHCO_3_, 1 mM NaH_2_PO_4_, 3 mM CaCl_2_, 4 mM MgCl_2_, 5 mM N-Tris (TES), pH 7.3. Odors at 10^-1^ dilution were delivered by switching mass-flow controlled carrier at 0.4 l/min and stimulus streams at 0.4 l/min (Sensirion) via software controlled solenoid valves (The Lee Company). This resulted in a final concentration of 5×10^-2^ of odor delivered to the fly. Air-streamed odor was delivered through a 1/16 inch ultra-chemical-resistant Versilon PVC tubing (Saint-Gobain, NJ, USA) that was placed 5 mm from the fly’s antenna.

Fluorescence was excited by a Ti-Sapphire laser (Mai Tai HP DS, 100 fs pulses) centered at 910 nm, attenuated by a Pockels cell (Conoptics) and coupled to a galvo-resonant scanner. Excitation light was focused by a 20X, 1.0 NA objective (Olympus XLUMPLFLN20XW), and emitted photons were detected by GaAsP photomultiplier tubes (Hamamatsu Photonics, H10770PA-40SEL), whose currents were amplified (Hamamatsu HC-130-INV) and transferred to the imaging computer (MScan 2.3.01). All imaging experiments were acquired at 30 Hz. When necessary, movies were motion-corrected using the TurboReg ^69^ ImageJ plugin ^70^. ΔF/F was calculated as was previously described^33^. When a pharmacological effect was tested in the same fly, all ROIs ΔF/F values were calculated with the baseline fluorescence of the ROI prior to pharmacology. We excluded non-responsive flies and flies whose motion could not be corrected.

#### Odors used

3-octanol (3-OCT) and 4-methylcyclohexanol (MCH) were purchased from Sigma-Aldrich (Rehovot, Israel) and were at the purest level available.

#### Optogenetic activation

For optogenetic experiments, brains were illuminated with 625 nm red light (GCS-0625-03; MIGHTEX LED) at 33 Hz (paired with a 5-s odor presentation).

#### Pharmacological application

The following drugs were used: Dopamine Hydrochloride (Alfa Aesar #A11136)), ACh (Sigma-Aldrich #A6625) and TTX (Alomone Labs #T-550). In all cases stock solutions were prepared and diluted in external solution to the final concentration before experiments. For local application, a glass pipette filled was placed in close proximity to the MB lobe and was emptied using a pico injector (Harvard Apparatus, PLI-100).

#### Structural Imaging

Brain dissections, fixation, and immunostaining were performed as described^71^. To visualize native GFP fluorescence, dissected brains were fixed in 4% (w/v) paraformaldehyde in PBS (1.86 mM NaH_2_PO_4_, 8.41 mM Na_2_HPO_4_, 175 mM NaCl) and fixed for 20 minutes at room temperature. Samples were washed for 3×20 minutes in PBS containing 0.3% (v/v) Triton-X-100 (PBS-T). Primary antisera were rabbit polyclonal anti-HA (1:500, ab9110, Abcam), and mouse monoclonal anti-Bruchpilot – nc82 (1:50, DSHB). Secondary antisera were Alexa488 coupled to goat anti-rabbit or Alexa647 coupled to goat anti-mouse (1:250, all Abcam). Primary antisera were applied for 2 days and secondary antisera for 1 day in PBS-T at 4 °C, followed by embedding in VECTASHIELD® PLUS Antifade Mounting Medium (H-1900, Vector laboratories). Images were collected on a Leica TCS SP5 or SP8 confocal microscope and processed in ImageJ.

#### RNA Purification, cDNA Synthesis, and Quantitative Real-time PCR Analysis

Total RNA from 60 adult heads was extracted using the EZ-RNA II kit (#20-410-100, Biological Industries, Kibbutz Beit-Haemek, Israel) for each biological replicate. Reverse transcription of total RNA (1000ng) into complementary DNA (cDNA) was performed using High-Capacity cDNA Reverse Transcription Kit with RNase Inhibitor (AB-4374966, Thermo Scientific, Massachusetts, USA). RT-PCR reactions were performed using Fast SYBR® Green Master Mix (AB-4385612, Applied Biosystems, CA, USA) in a StepOnePlus instrument (Applied Biosystems, CA, USA). Primers (β-Tubulin, forward primer, CCAAGGGTCATTACACAGAGG, reverse primer, ATCAGCAGGGTTCCCATACC; mAChR-B, forward primer, ATGCGGTCGCTTAACAAGTC, reverse primer, GCTCCCTTCTAAGGCTCCAG) were calibrated, and a negative control was performed for each primer. Samples were measured in technical triplicates, and values were normalized according to mRNA levels of the β-Tubulin housekeeping gene. The amplification cycles were 95°C for 30s, 60°C for 15s, and 72°C for 10s. At the end of the assay, a melting curve was constructed to evaluate the specificity of the reaction. The fold change for each target was subsequently calculated by comparing it to the normalized value of the ELAV-gal4 parent. Quantification was assessed at the logarithmic phase of the PCR reaction using the 2-ΔΔCT method, as described previously ^72^.

#### Connectome Data Analysis

For the connectome analysis, the Hemibrain v1.2.1 dataset made publicly available by Janelia Research Campus ^41,73^ was used. Hemibrain data was accessed via the Neuprint python package (https://github.com/connectome-neuprint/neuprint-python). To distinguish between axo-axonal connections and dendro-dendritic connections, the connections were filtered based on the ROI in which they occur (i.e. Calyx for dendrites or α/β, α’/β’ or √lobe).

### QUANTIFICATION AND STATISTICAL ANALYSIS

#### Statistics

Statistical analyses were carried out in GraphPad Prism as described in figure legends and **Table S1**.

**Table S1**: Statistical analysis. Related to Figures 2, 3, 4, 5, 6, 7, S1, S2, S3, S4, S5, and S6.

---

**Figure S1:**
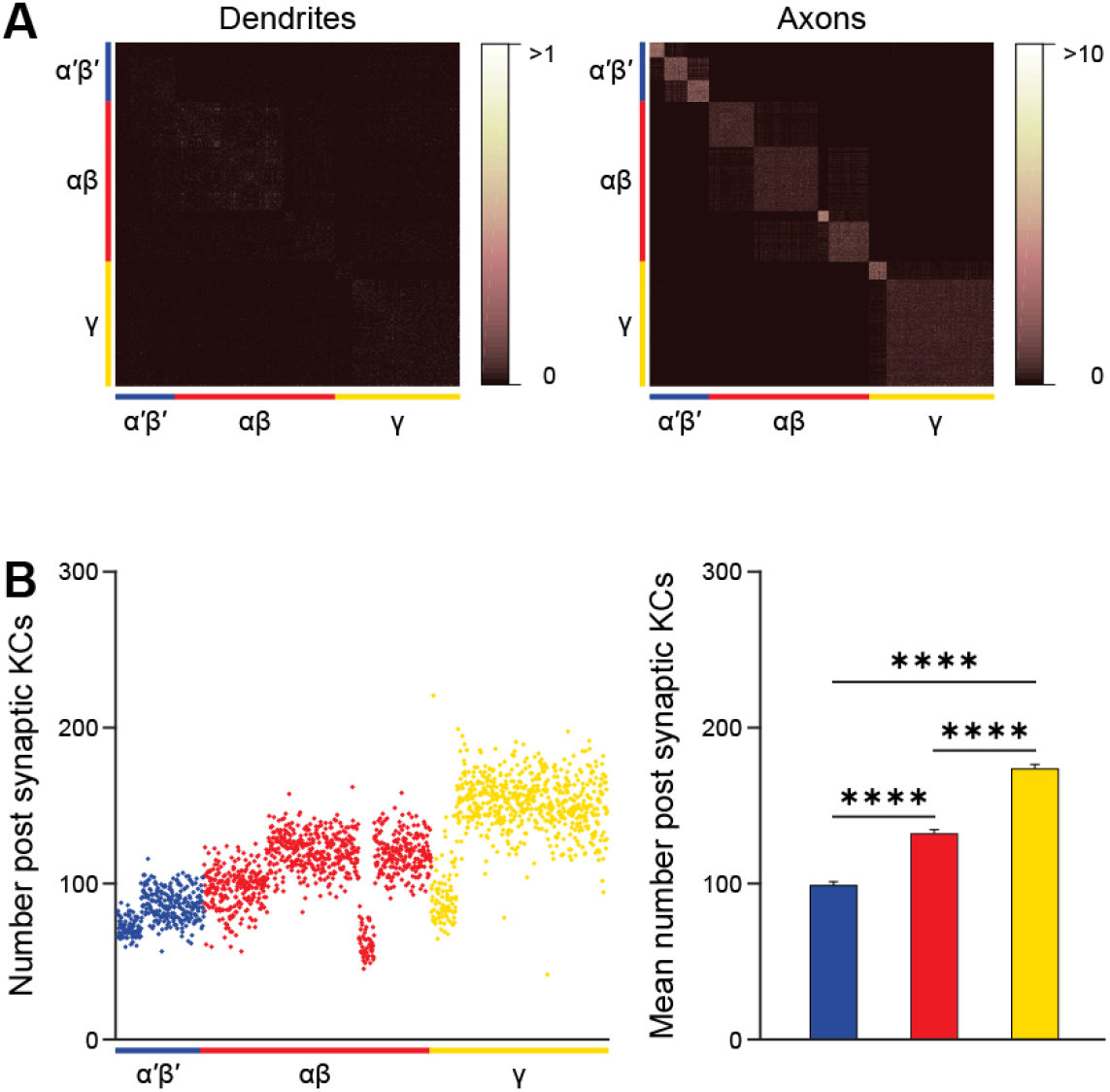
γ KCs have strong lateral axo-axonal connections. Related to Figure 1. **A.** Dendritic (left) and axonal (right) lateral KC-KC connections, note different scale. The number of synapses made between each KC to all other KCs, arranged by the three main subclasses of KCs, in the different regions of the MB (Calyx and Lobes) are shown. KCs show strong axonal connections to other cognate KCs. Blue (α’β’), red (αβ) and yellow (γ), indicate the KC subtypes as designated. **B.** The number of post-synaptic KCs each KC has according to the different types of KCs. Blue, red and yellow, indicate the KC subtypes as designated. **C.** Mean number of post-synaptic KCs, obtained from the data presented in B. Blue, red and yellow, indicate the KC subtypes as designated. γ KCs have the highest number of post-synaptic KC partners (mean ± SEM), n (left to right): 337, 889, 689; **** p<0.0001; (Kruskal-Wallis Dunn’s correction for multiple comparisons. For detailed statistical analysis see **Table S1**.

**Figure S2:**
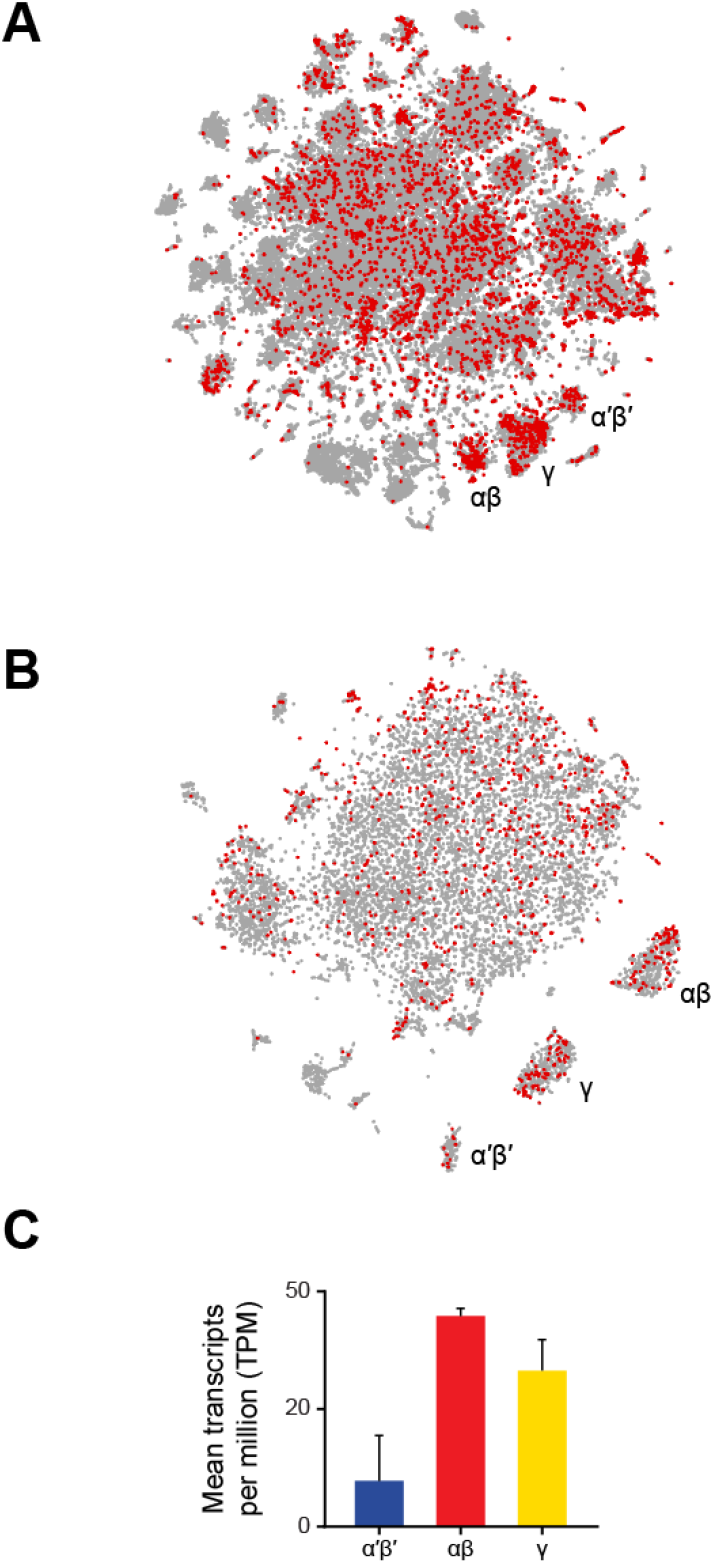
Expression of mAChR-B from single-cell transcriptome profiling. Related to Figure 1. **A.** Data from Davie et al., 2018. 56,902 *Drosophila* brain cells arranged according to their single-cell transcriptome profiles, along the top 2 principal components using t-SNE. Red coloring indicates expression of mAChR-B. KC subtype clusters are labeled as identified in Davie et al., 2018. **B.** As in A but with data from Croset et al., 2018 (10,286 *Drosophila* brain cells). **C.** Data from Aso et al. 2019. (2500 γ and αβ KCs, 1000 α’β’ KCs). Blue (α’β’), red (αβ) and yellow (γ), indicate the KC subtypes as designated. For A, B, Images screenshotted from SCope (http://scope.aertslab.org) on 9 March 2022.

**Figure S3:**
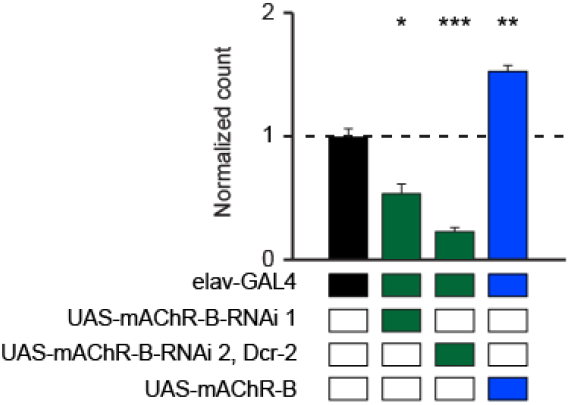
mAChR-B RNAi and overexpression efficiency. Related to Figure 2. qRT-PCR of mAChR-B with both UAS-mAChR-B RNAi (RNAi 1 and 2) and with UAS-mAChR-B driven by elav-GAL4. The housekeeping gene β-Tubulin was used for normalization. Knockdown flies have ~55% and 25% for RNAi 1 and 2 respectively and overexpression of mAChR-B has ~155% of the control levels of mAChR-B mRNA (mean ± SEM; 3 biological replicates each with 3 technical replicates; * p < 0.05; (Multiple T-test with Holm-Šídák correction for multiple comparisons). For detailed statistical analysis see **Table S1**.

**Figure S4:**
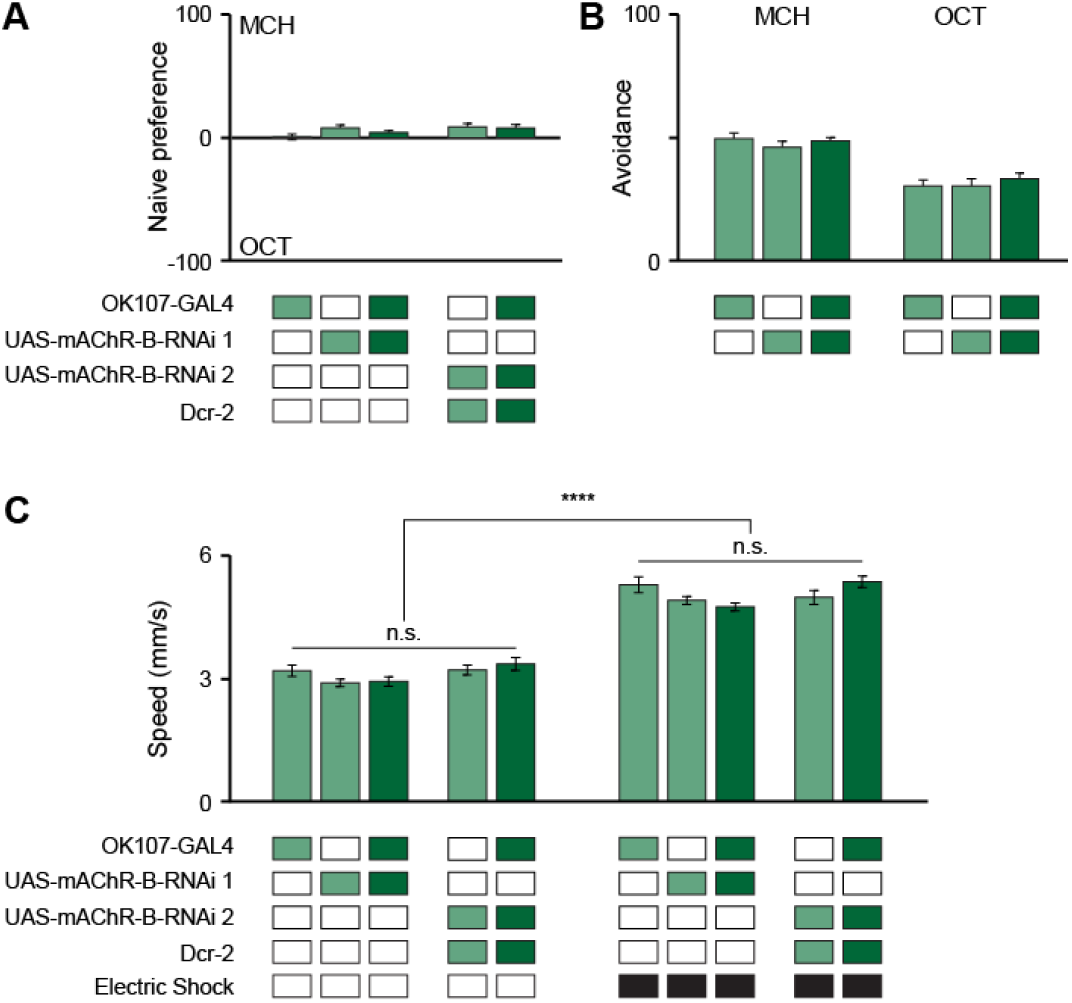
Control experiments for Figure 2. Related to Figure 2. **A.** mAChR-B KD flies show normal preference between OCT and MCH compared to their genotypic controls (mean ± SEM), n (left to right): 56, 162, 77, 66, 61 (Welch and Brown-Forsythe tests-Dunnett’s multiple comparisons test). For detailed statistical analysis see **Table S1**. **B.** mAChR-B KD flies show normal olfactory avoidance of OCT and MCH compared to their genotypic controls (mean ± SEM), n (left to right): MCH: 96, 96, 88; OCT: 108, 75, 95 (Kruskal-Wallis Dunn’s correction for multiple comparisons). For detailed statistical analysis see **Table S1**. **C.** Sensitivity to shock (extent to which flies walk faster while being shocked) is not affected by knocking down mAChR-B in KCs. Walking speed with (right) or without (left) an electric shock is presented. mAChR-B KD did not affect walking speed in either condition (mean ± SEM, n (left to right): no shock: 49, 162, 77, 66, 61; with shock: 49, 162, 77, 66, 61 (Kruskal-Wallis Dunn’s correction for multiple comparisons). For detailed statistical analysis see **Table S1**.

**Figure S5:**
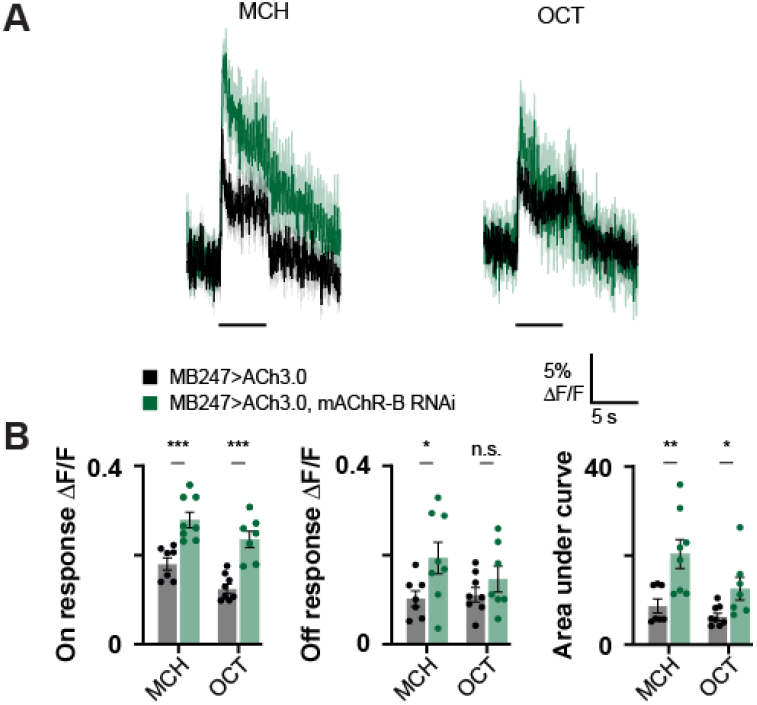
mAChR-B KD increase γ KC synaptic release. Related to Figure 3. Synaptic release as indicated by ACh signal in KCs following MCH or OCT was measured in control flies (MB247-GAL4>UAS-GACh3.0) and KD flies (MB247-GAL4>UAS-GACh3.0, UAS-mAChR-B). **A.** ΔF/F of GACh3.0 signal in the lobe of γ KCs for control (black) and KD (green) flies, during presentation of odor pulses (horizontal lines). Data are mean (solid line) ± SEM (shaded area). **B.** Peak “on” response (left), Peak “off” response (middle), and the integral of the odor response (right) of the traces presented in A (mean ± SEM). n for control and KD flies, respectively: MCH,7, 8; OCT, 8, 7. * p < 0.05, ** p < 0.01, (Mann-Whitney test with Holm Šídák correction for multiple comparisons). For detailed statistical analysis see **Table S1**.

**Figure S6:**
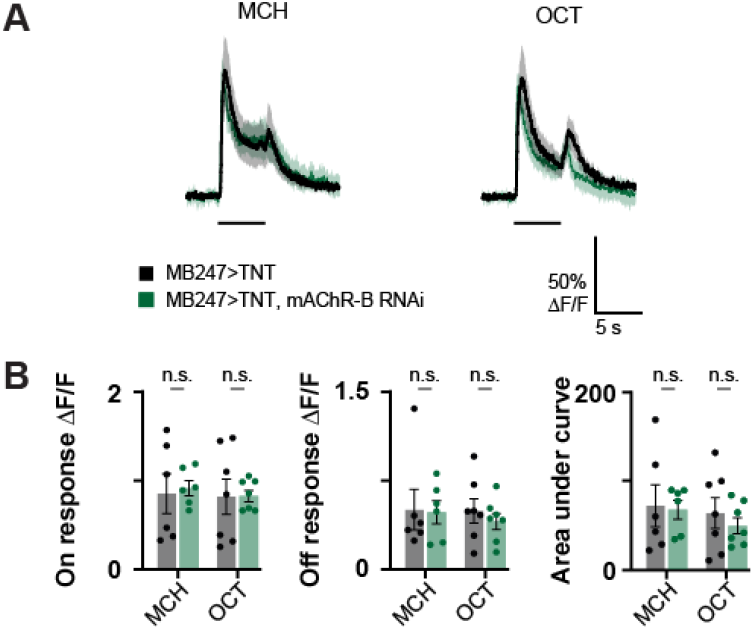
mAChR-B effect is post-synaptic to KC release. Related to Figure 6. KC odor responses to MCH and OCT were measured in control flies (MB247-GAL4>UAS-GCaMP6f) and knockdown flies (MB247-GAL4>UAS-GCaMP6f, UAS-mAChR-B RNAi 1) when KC synaptic release was blocked using UAS-TNT. **A.** ΔF/F of GCaMP6f signal in the lobe of γ KCs for control (black) and KD (green) flies, during presentation of odor pulses (horizontal lines). Data are mean (solid line) ± SEM (shaded area). **B.** Peak “on” response (left), Peak “off” response (middle), and the integral of the odor response (right) of the traces presented in A (mean ± SEM). n for control and KD flies, respectively: MCH, 6, 6; OCT, 7, 7. No statistical difference is observed between control and mAChR-KD flies (Mann-Whitney test with Holm Šídák correction for multiple comparisons). For detailed statistical analysis see **Table S1**.

**Figure S7:**
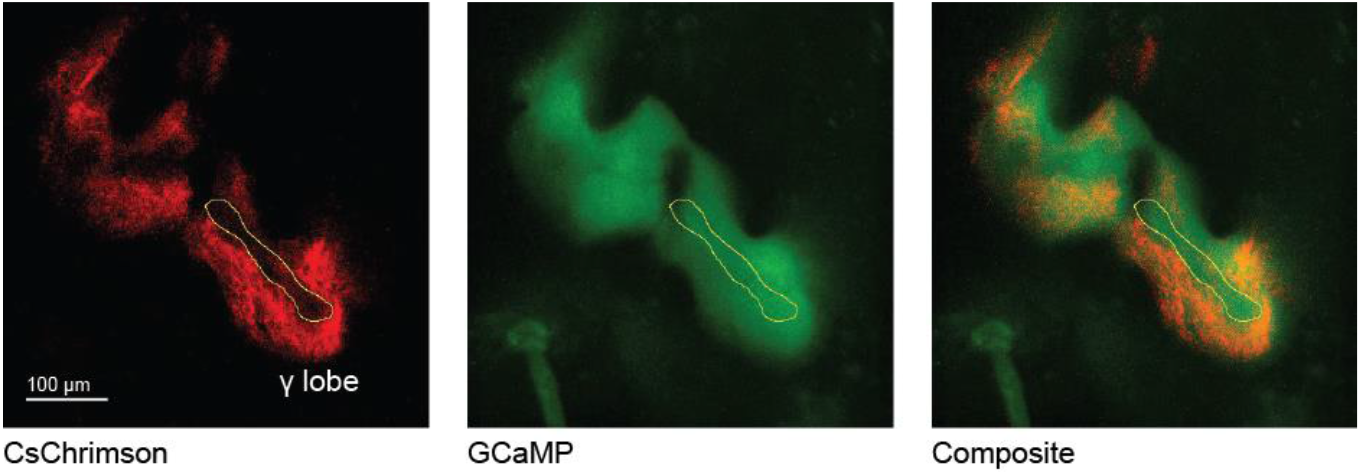
2-photon analysis for Figure 6. Related to Figure 6. Example of region selection for the analysis presented in Figure 5. A single plane average intensity projection over time (500 frames) of a 2-photon image obtained from a fly carrying MB247-LexA-LexAop-GCaMP6f, and CsChrimson::tdTomato in stochastically distributed subsets of neurons using TI{20XUAS-SPARC2-I-Syn21-CsChrimson::tdTomato-3.1}CR-P40 within the MB247-GAL4 driver line transgenes. ROIs were selected manually in Fiji to include only GCaMP labeled areas and not tdTomato. *Left*, CsChrimson is only partially expressed in the γ lobe. *Middle*, GCaMP6f signal throughout the γ lobe. *Right*, a composite of the CsChrimson and GCaMP6f signals.

**Figure S8:**
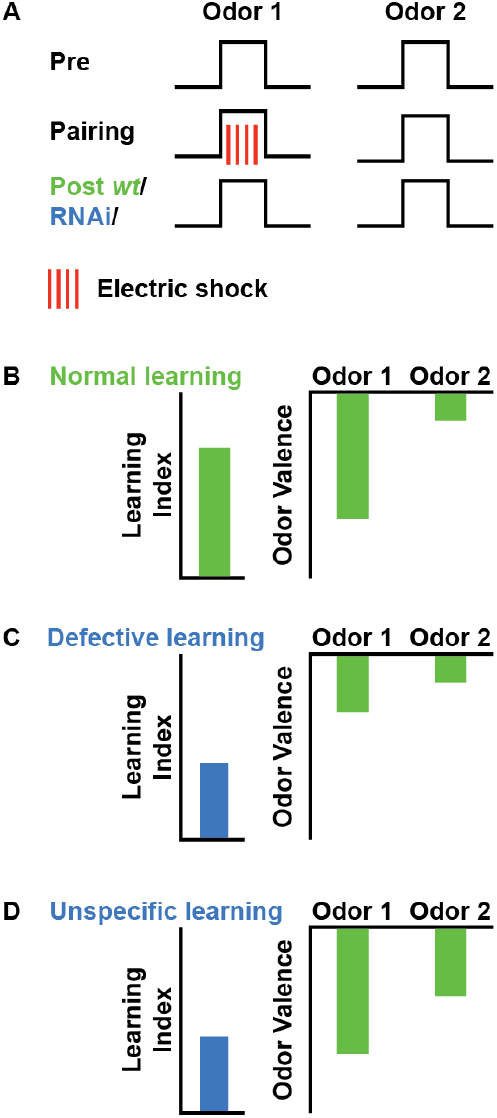
Comparison between defective and unspecific learning. Related to Figure 7. **A.** Illustration of the classical conditioning protocol. The preference between odor 1 and odor 2 is evaluated prior to conditioning. Odors are then subjected to a conditioning protocol in which odor 1 (CS^+^) is associated with an electric shock. This is then followed by another examination of odor preference. **B.** Under normal conditions following conditioning, the valence of odor 1 (CS^+^) becomes very negative whereas that of odor 2 (CS^-^) is not affected. **C.** When defective conditioning occurs, for example in the case where the dopaminergic neurons are inactive or when the dopaminergic receptors on KCs are knocked down, odor 1 (CS^+^) valence is not as negative as under normal conditions. Thus, the difference between the valence of odor 1 and odor 2 becomes smaller, and the learning index is reduced. **D.** When unspecific conditioning occurs, as suggested following mAChR-B KD, the valence of odor 1 (CS^+^) becomes very negative, as under normal conditions. However, the valence of odor 2 (CS^-^) is also affected even if to a lesser extent. Thus, the difference between the valence of odor 1 and odor 2 becomes smaller and the learning index is reduced, in a similar manner to defective learning. The underlying mechanism, however, is different.

**Figure S9:**
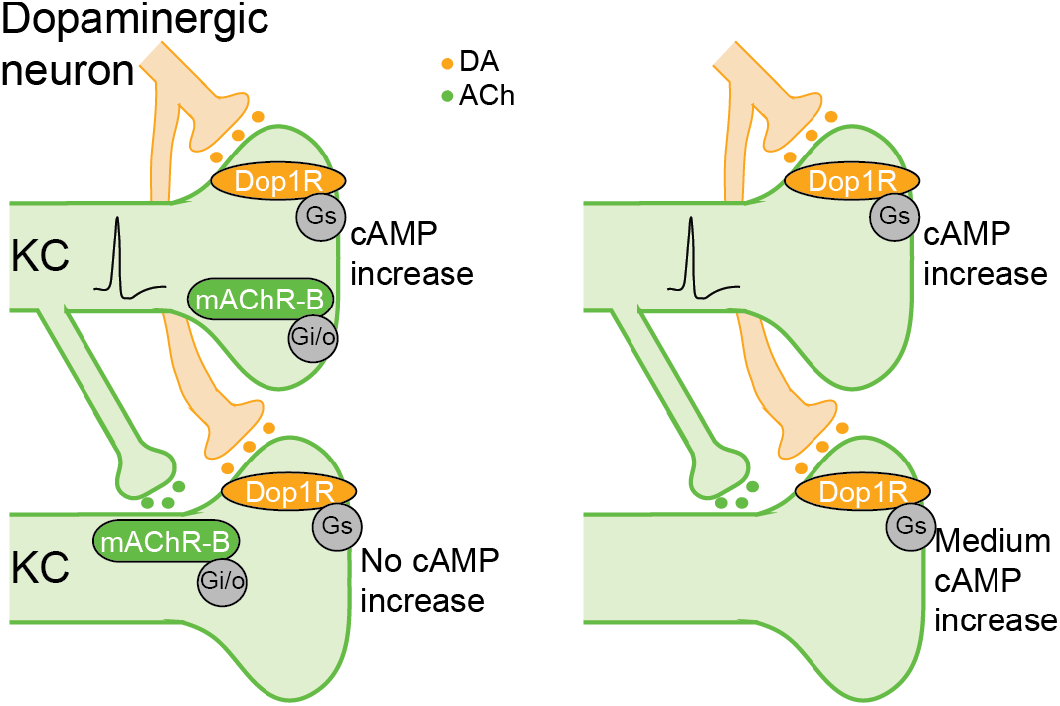
A model of mAChR-B lateral neuromodulation and noise free learning. Related to Figure 7. *Left*, under normal conditions when pairing of an electric shock with an odor occurs, the CS^+^ activates a subset of KCs and DA is released on all KCs. DA coincidence with KC activity results in a large increase in cAMP (due to Ca^2+^ increase in KC presynaptic terminal, which is required for maximal activity of adenylate cyclase, top) and, as a result, induction of plasticity. DA also activates dopaminergic receptors on the less or non-active KCs (bottom). The active KCs release ACh, activating mAChR-B of their cognate KCs. This mAChR-B neuromodulation reduces cAMP and directly opposes the DA neuromodulation, resulting in suppression of cAMP increase in non-active KCs (bottom). In addition, mAChR-B decreases the Ca^2+^ elevation in KC presynaptic terminals. In the case of KCs that are non-active and weakly activate (and are therefore not the main carrier of the CS+ odor signal), this cholinergic neuromodulation will prevent DA neuromodulation (bottom). *Right*, when mAChR-B is KD, the high cAMP increase following DA in active KCs is not affected (top). However, in non-active and weakly active KCs there is no mAChR-B to counter the cAMP increase caused by DA. As a result, there is an increase in cAMP, even if to a lesser extent than that which occurs in active KCs. As a consequence, some plasticity occurs also in the off-target KCs. These KCs naturally do not respond reliably to the conditioned odor but rather to other odors. Thus, unspecific plasticity and conditioning can occur.

**Table S1:**
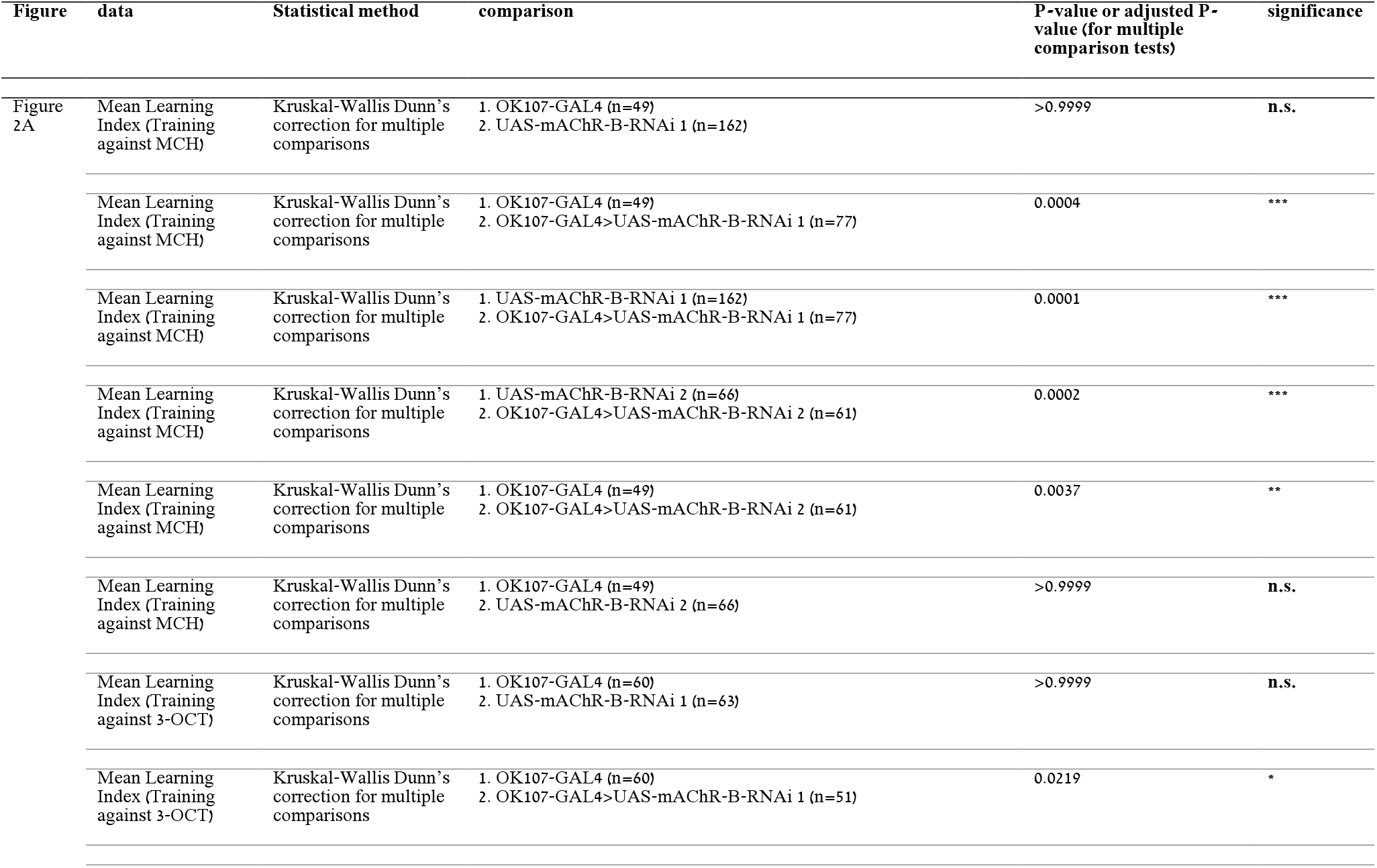
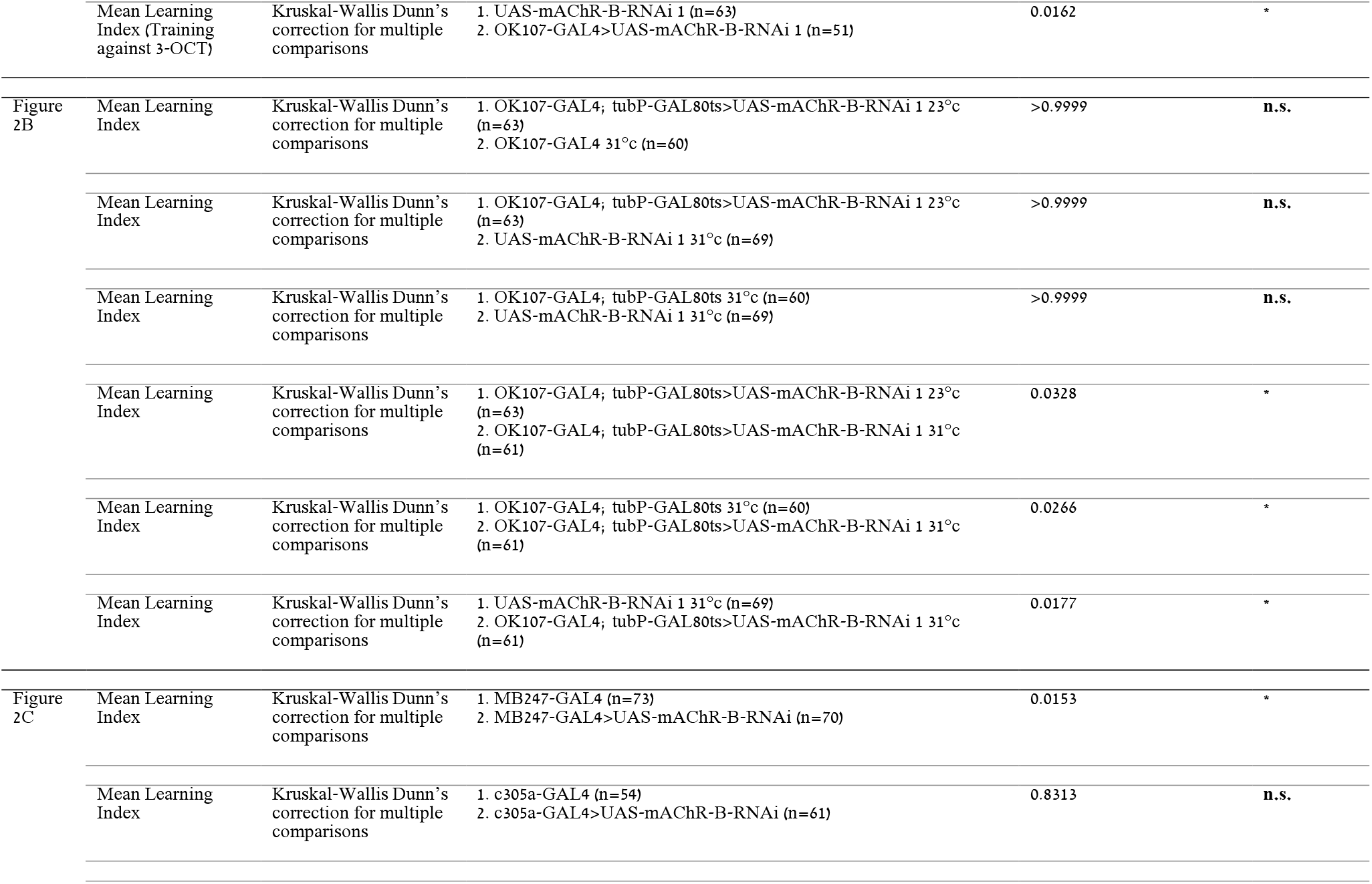
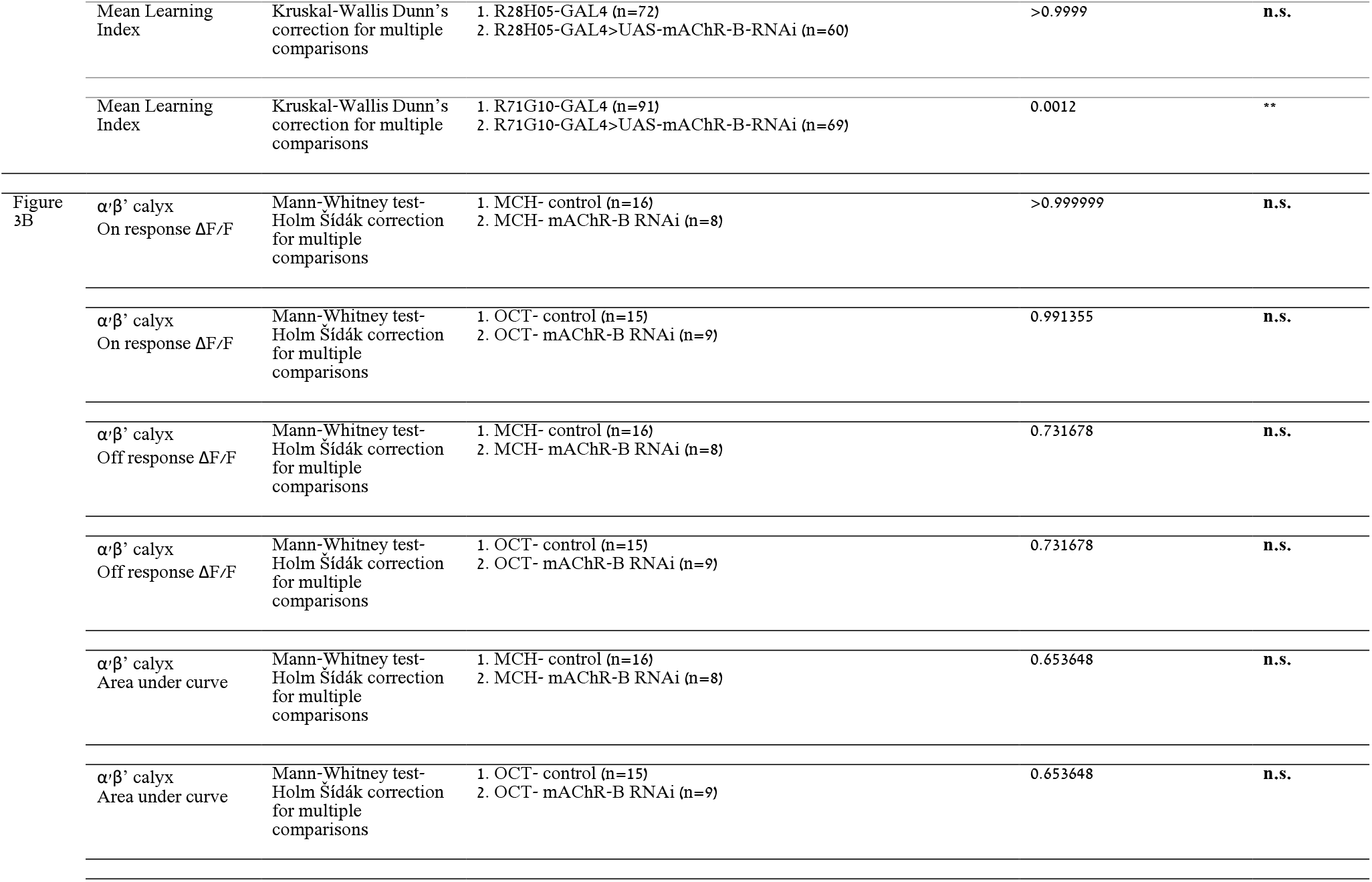
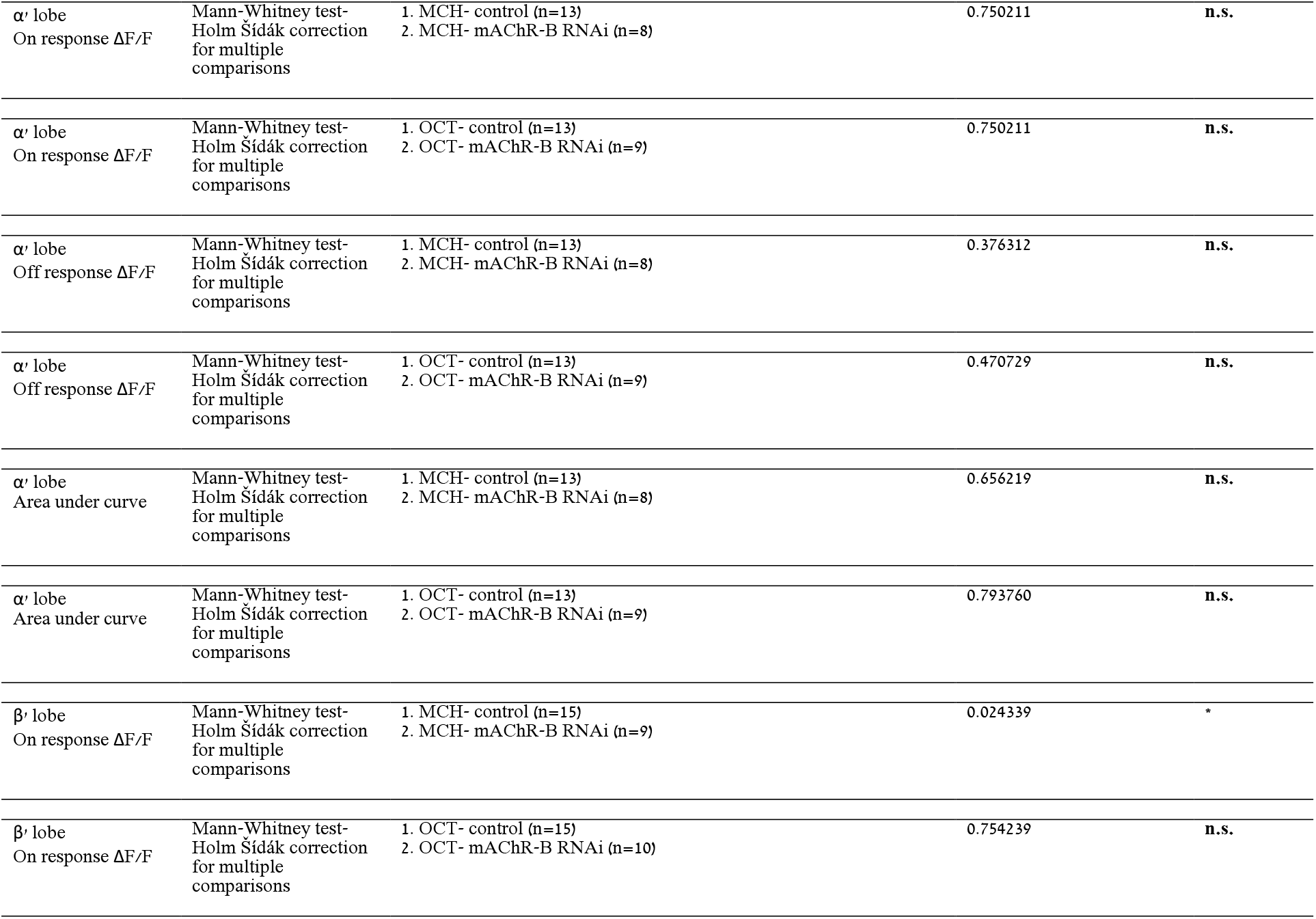
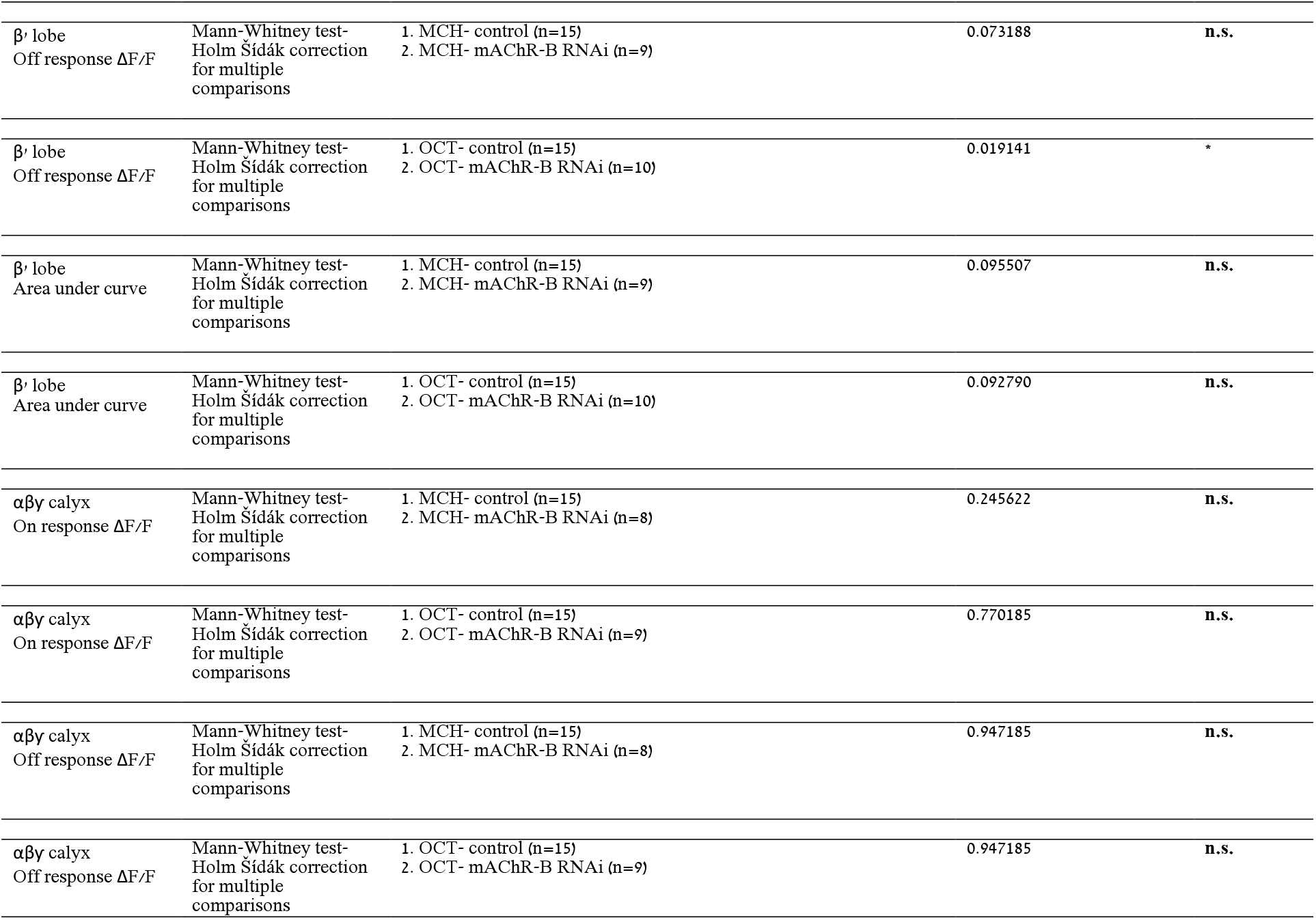
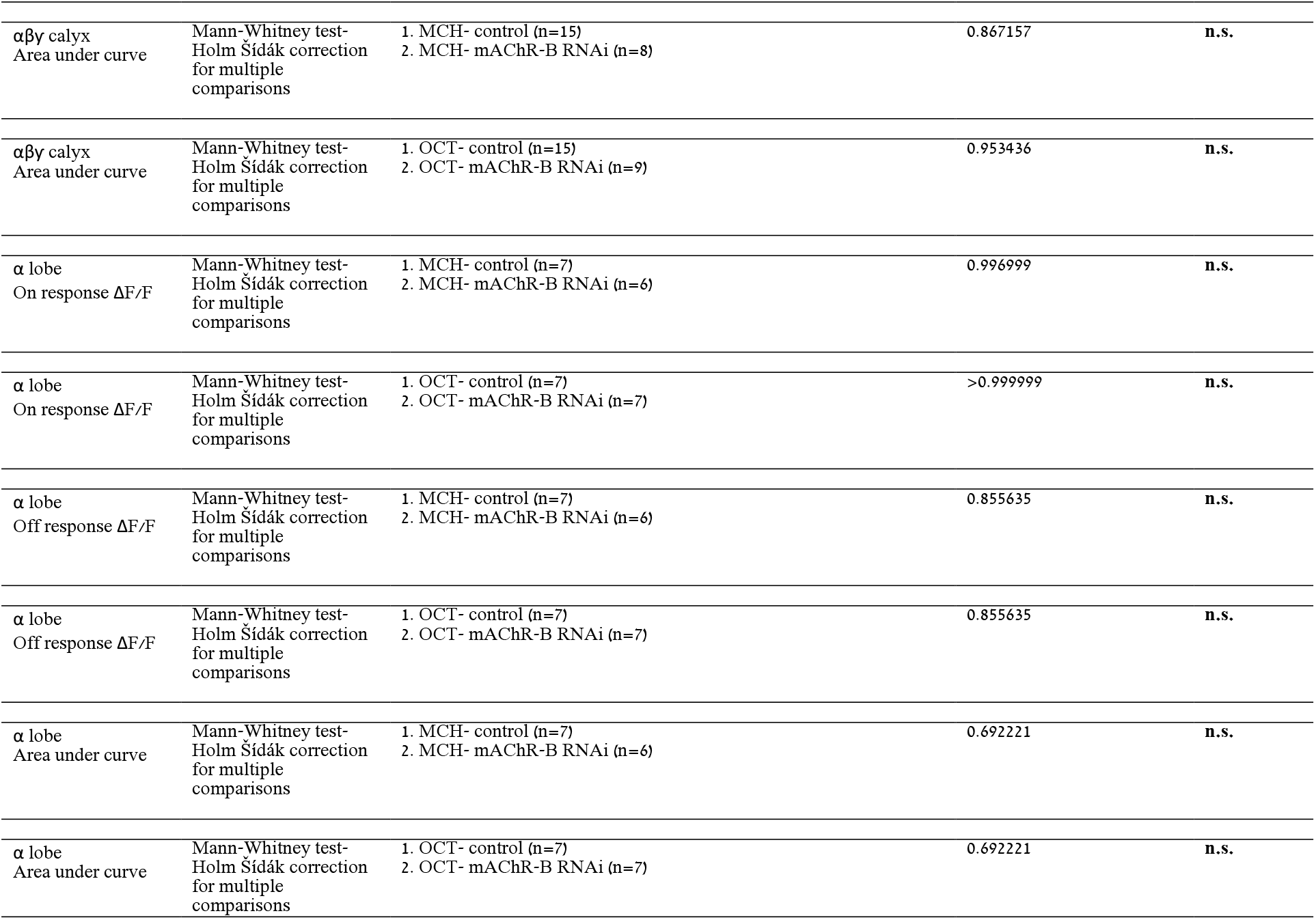
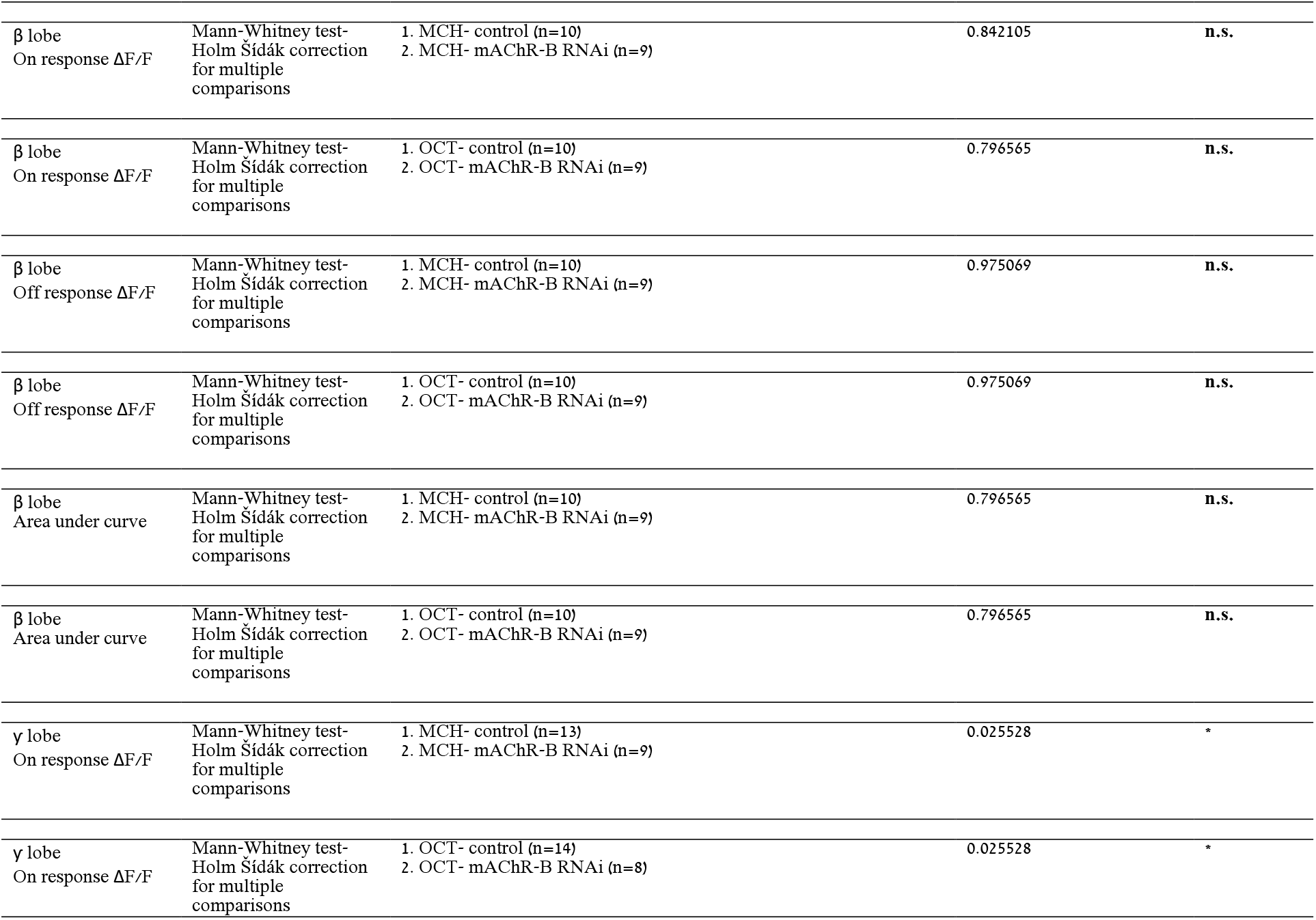
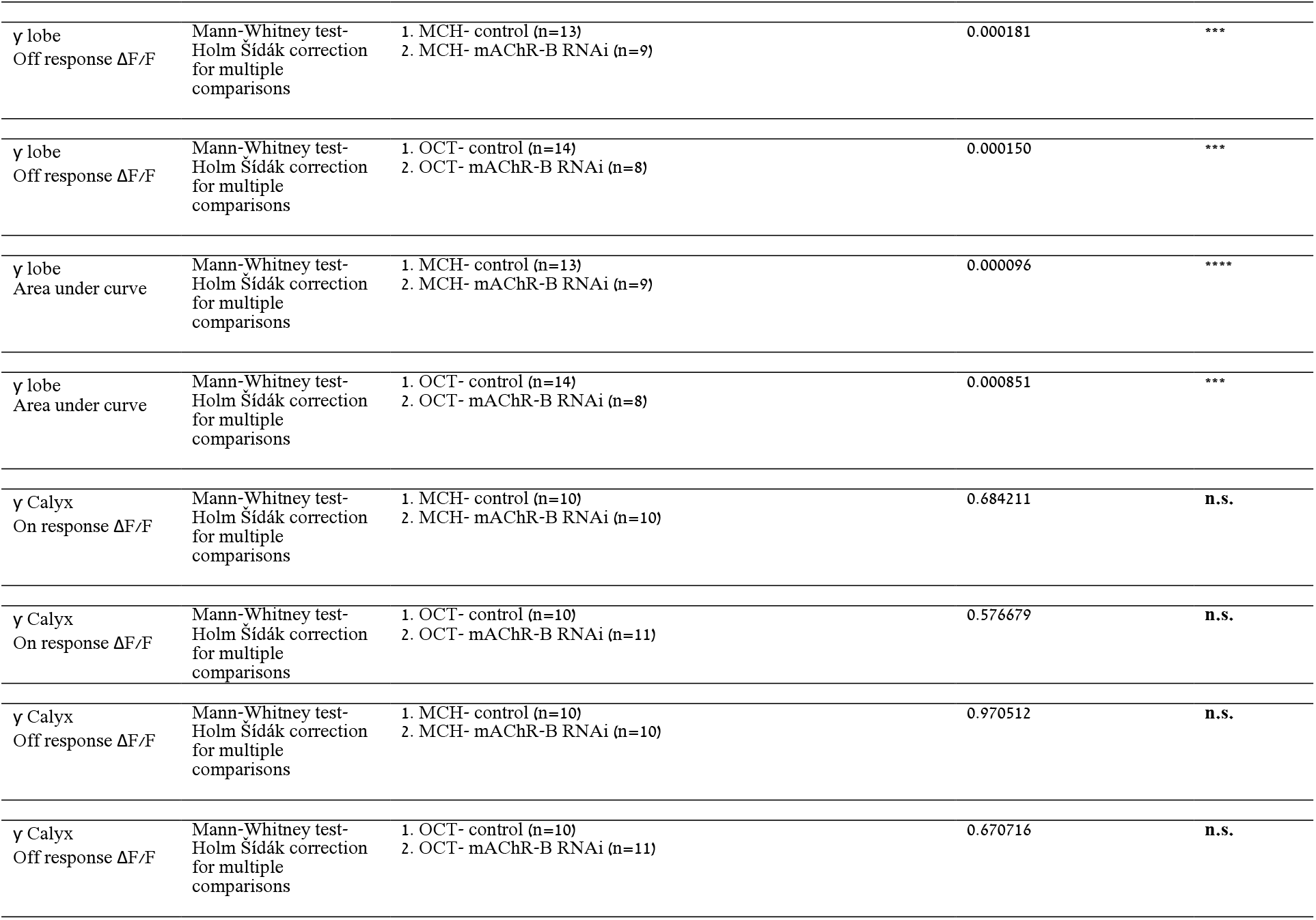
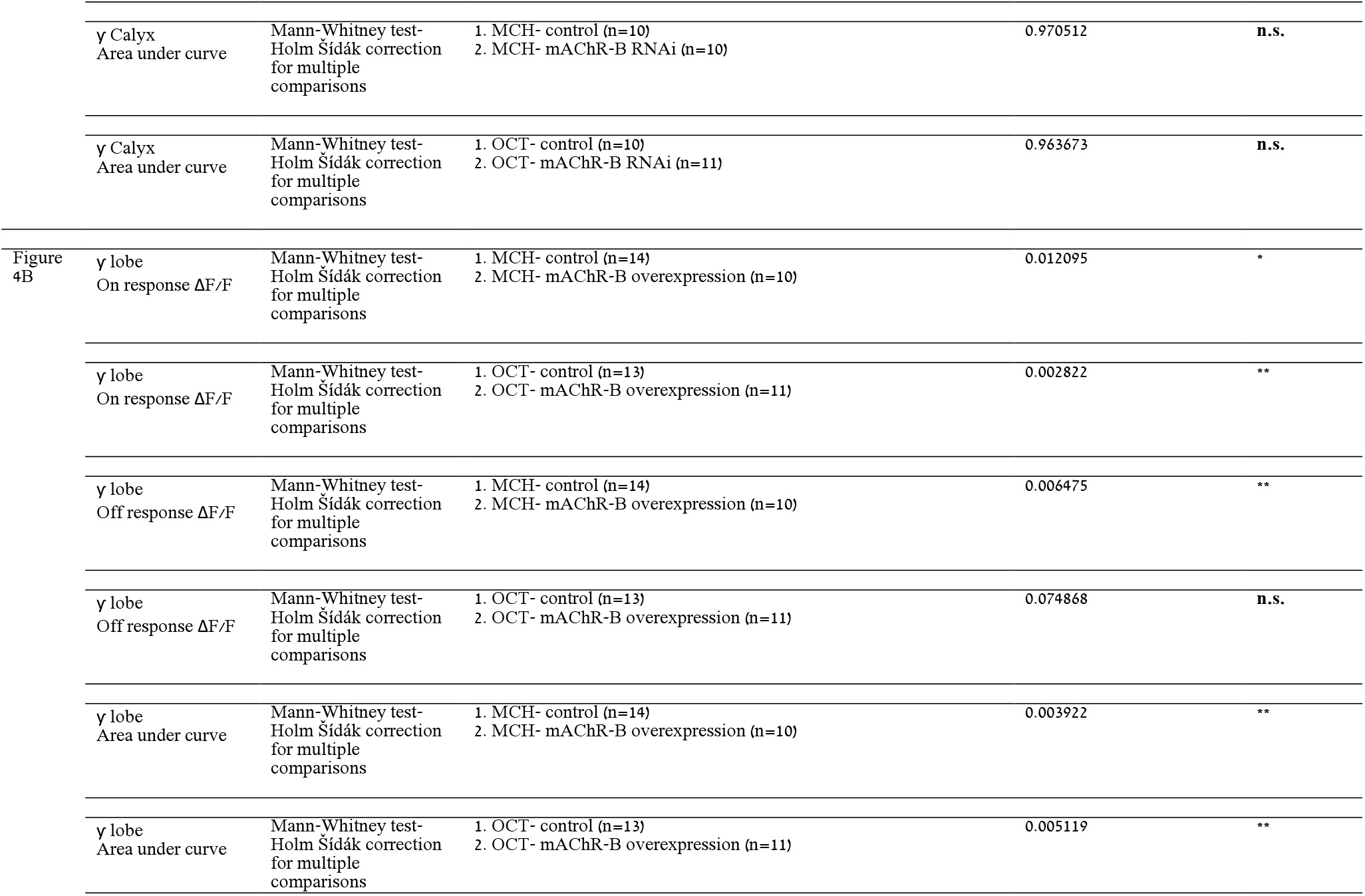
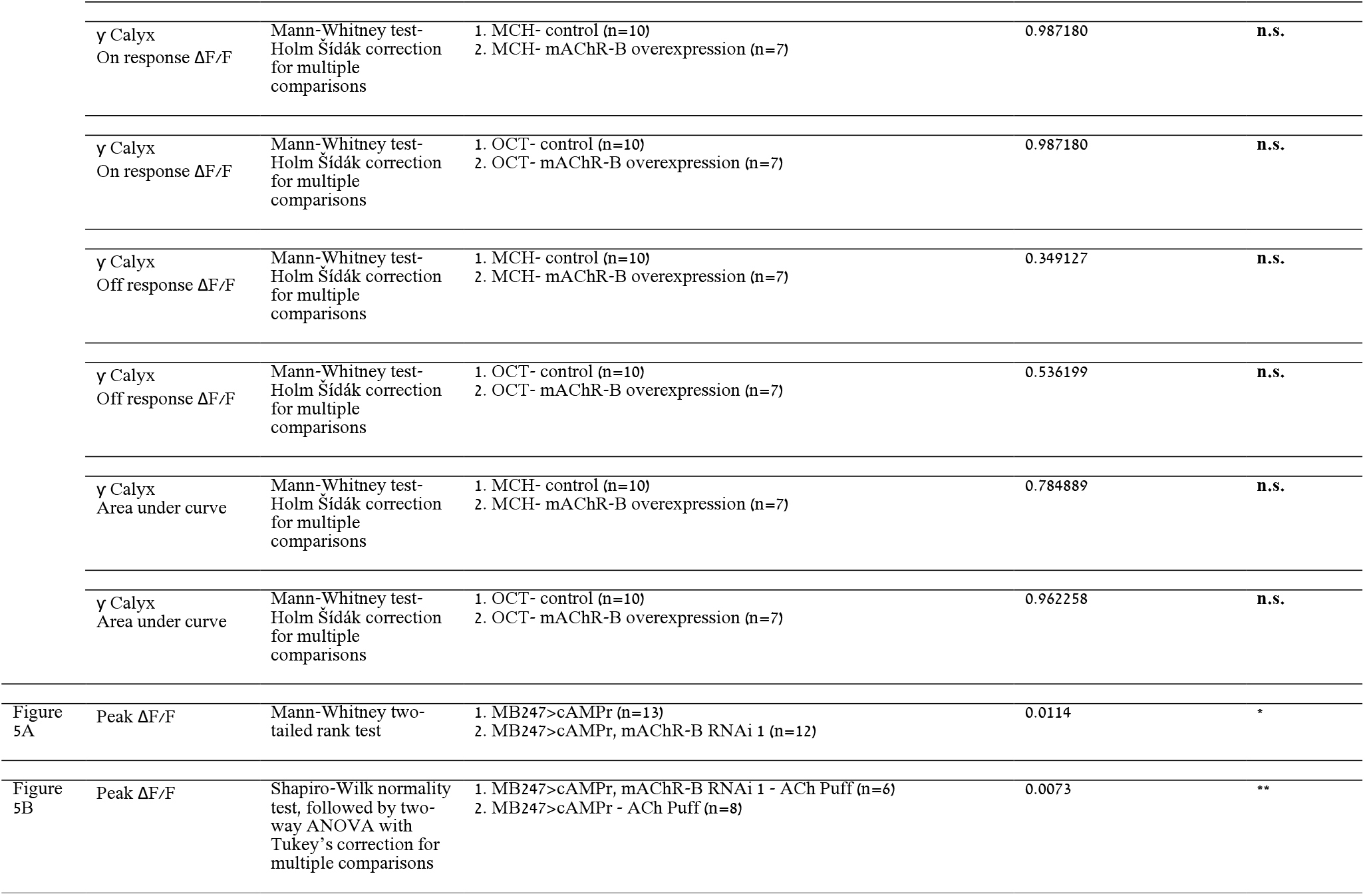
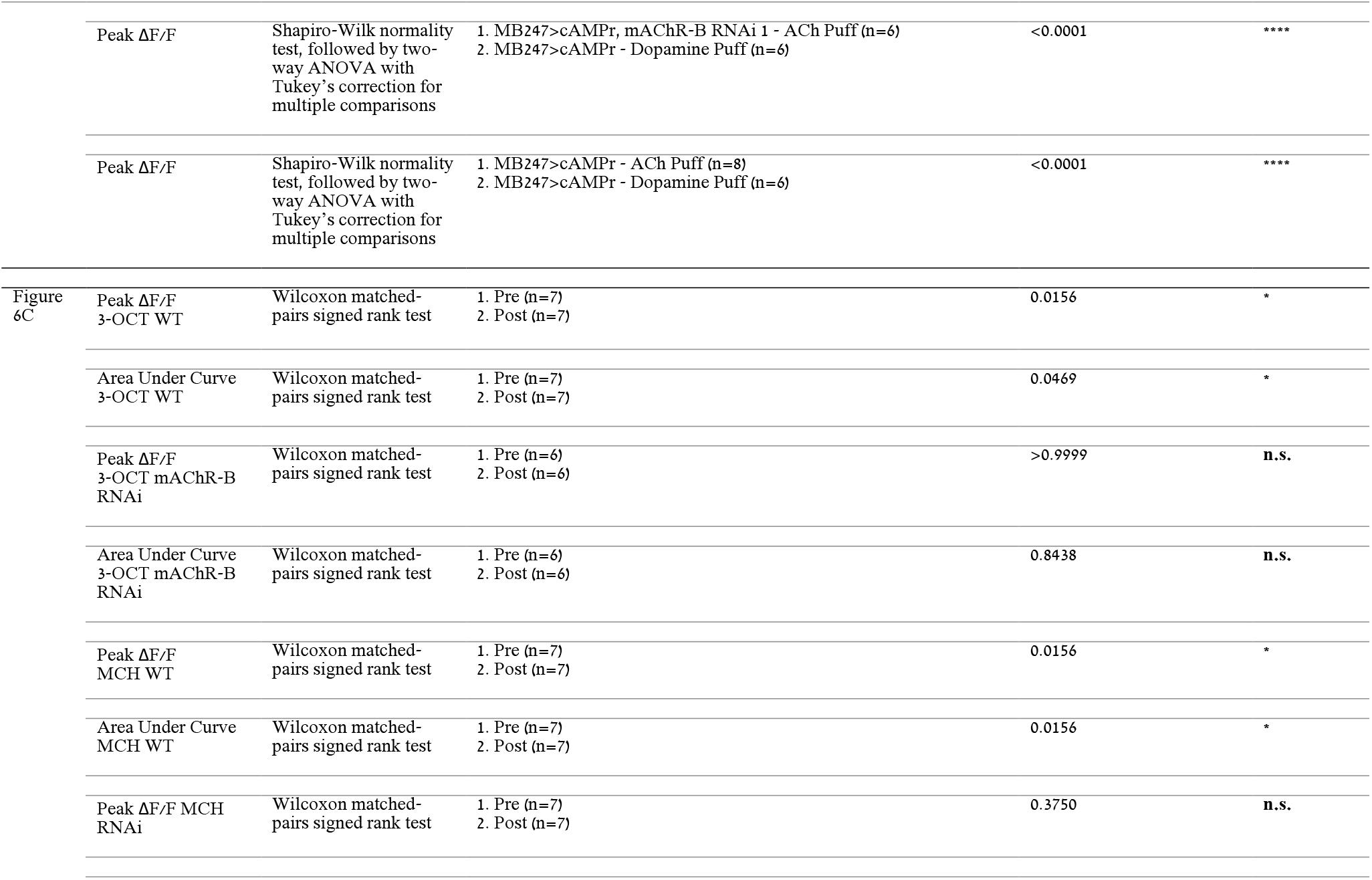
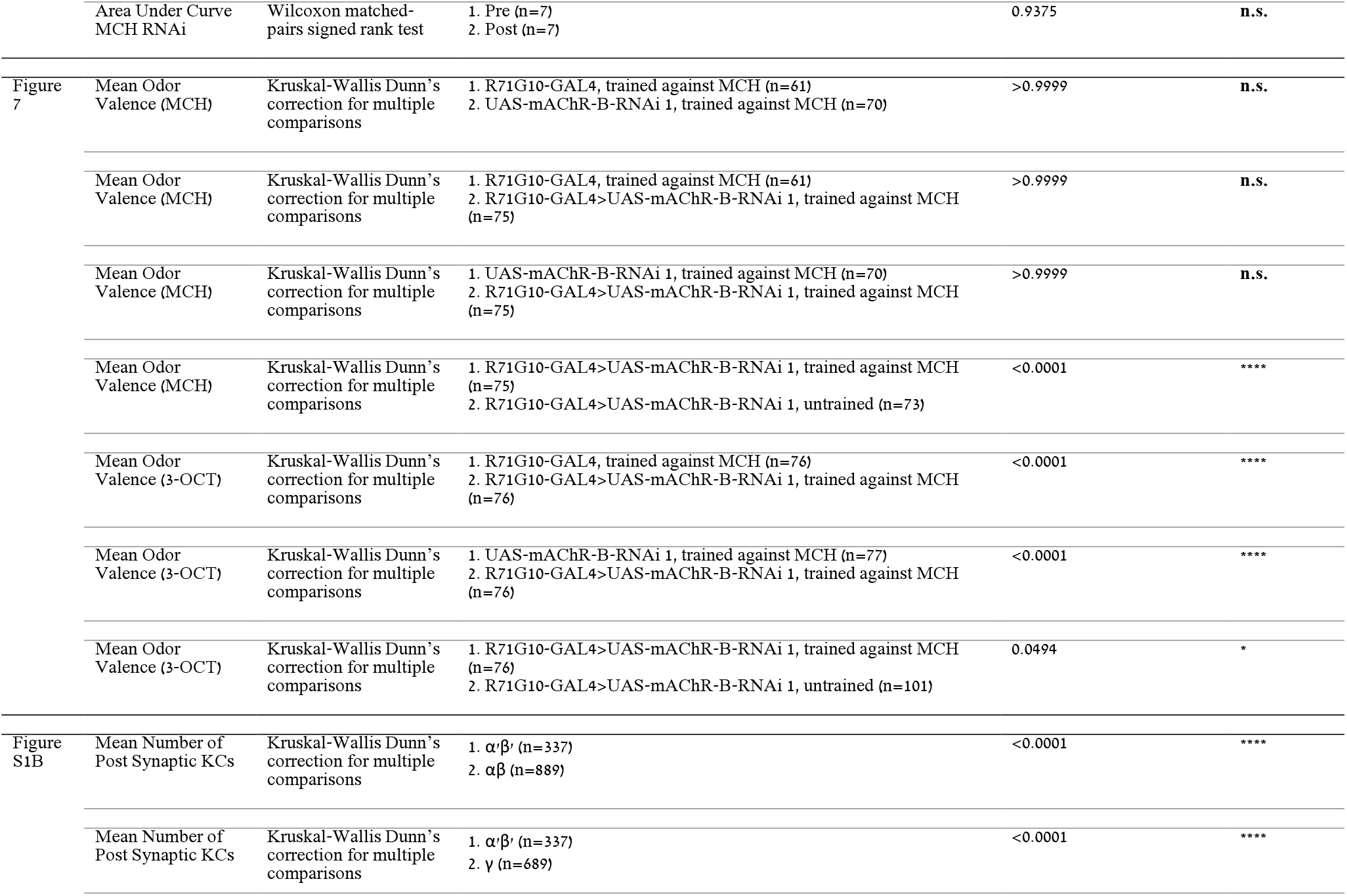
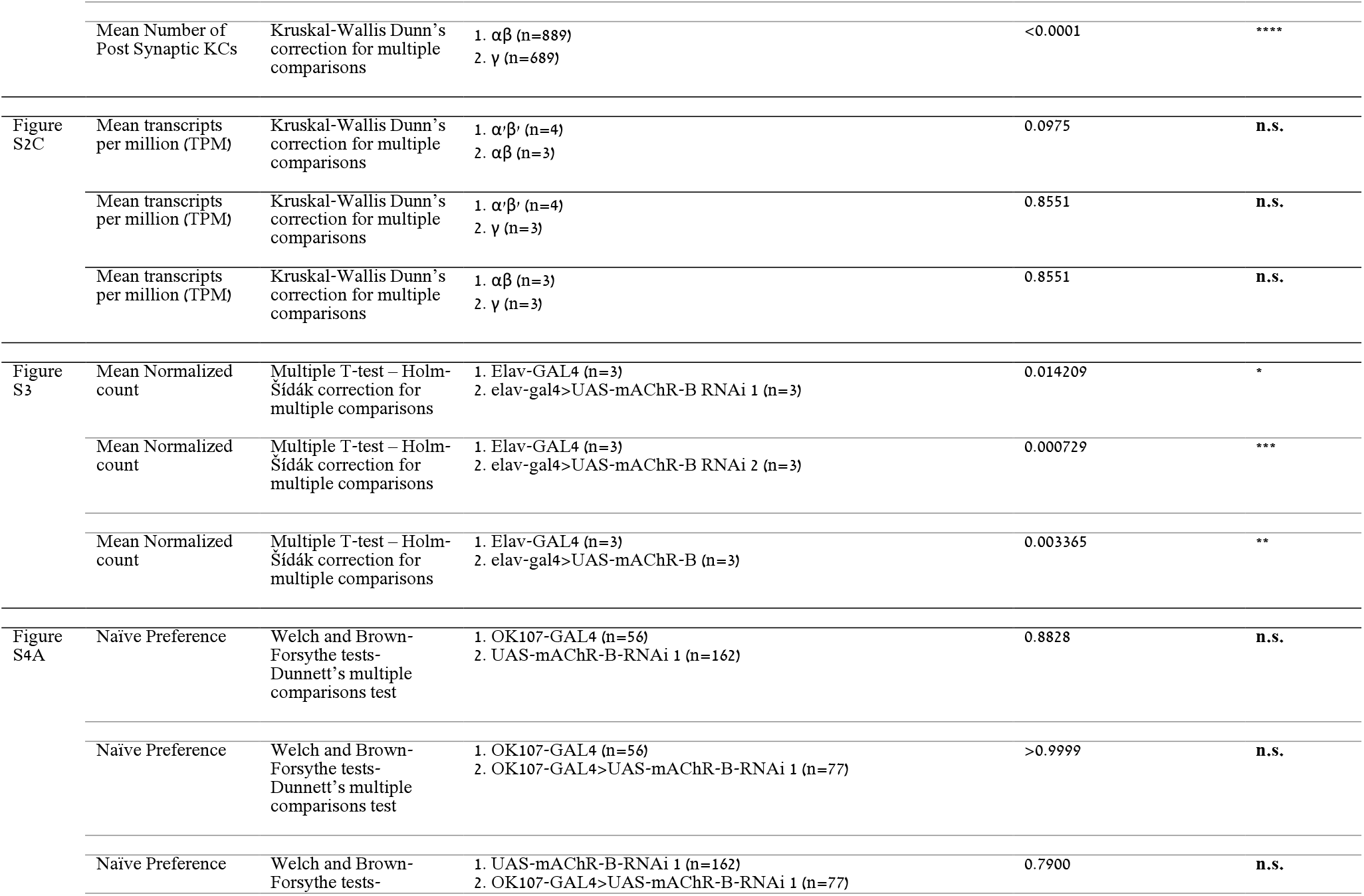
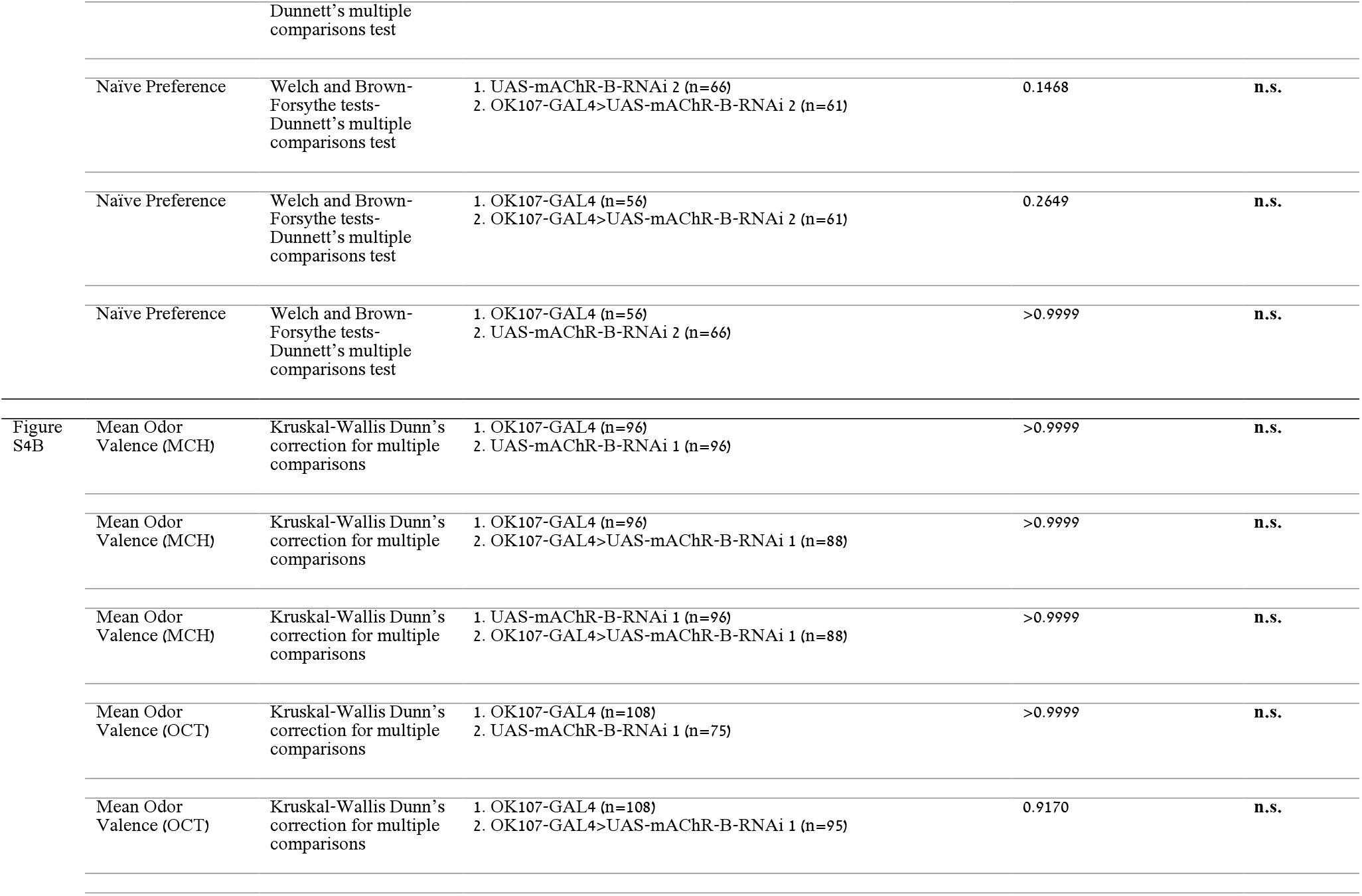
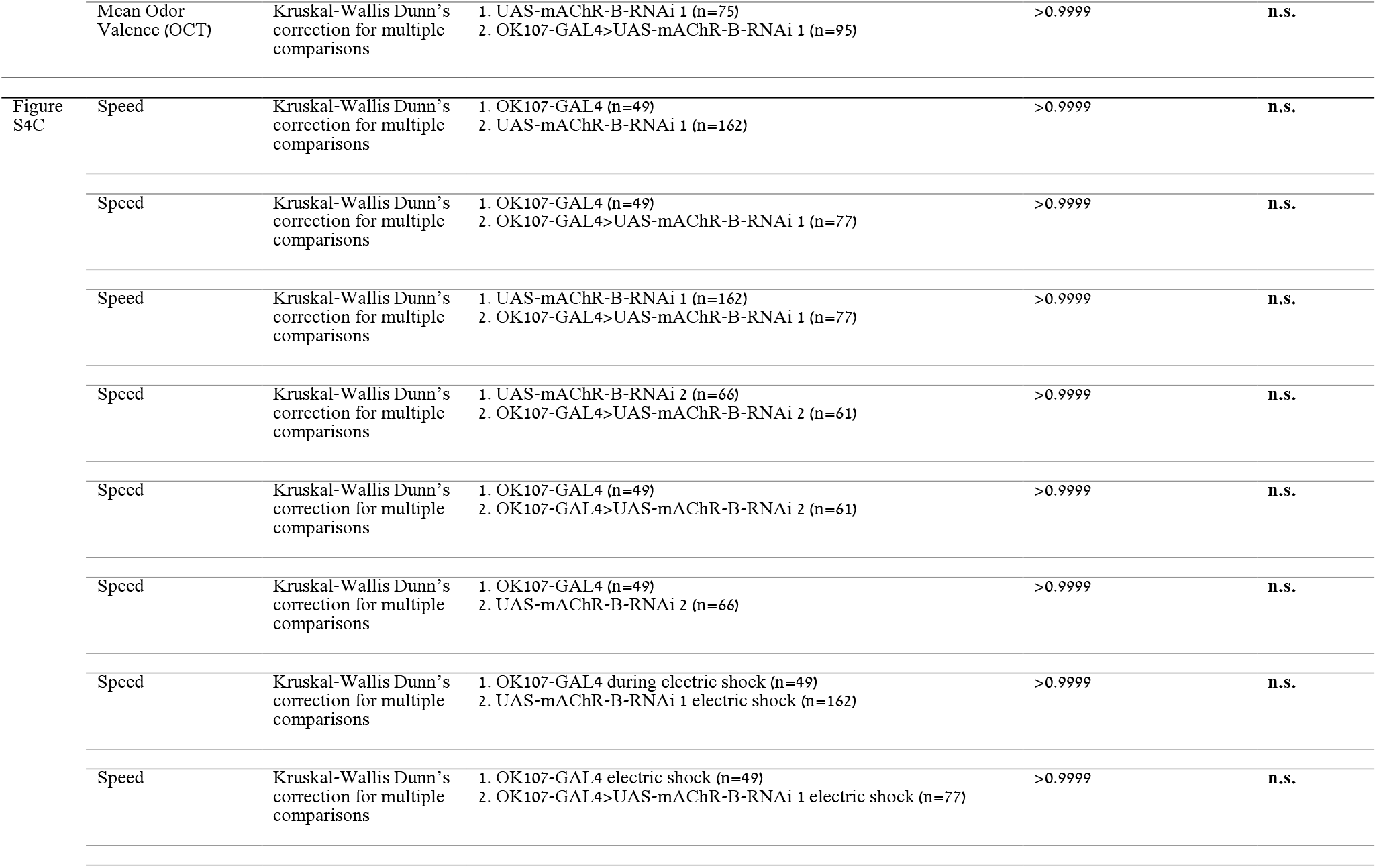
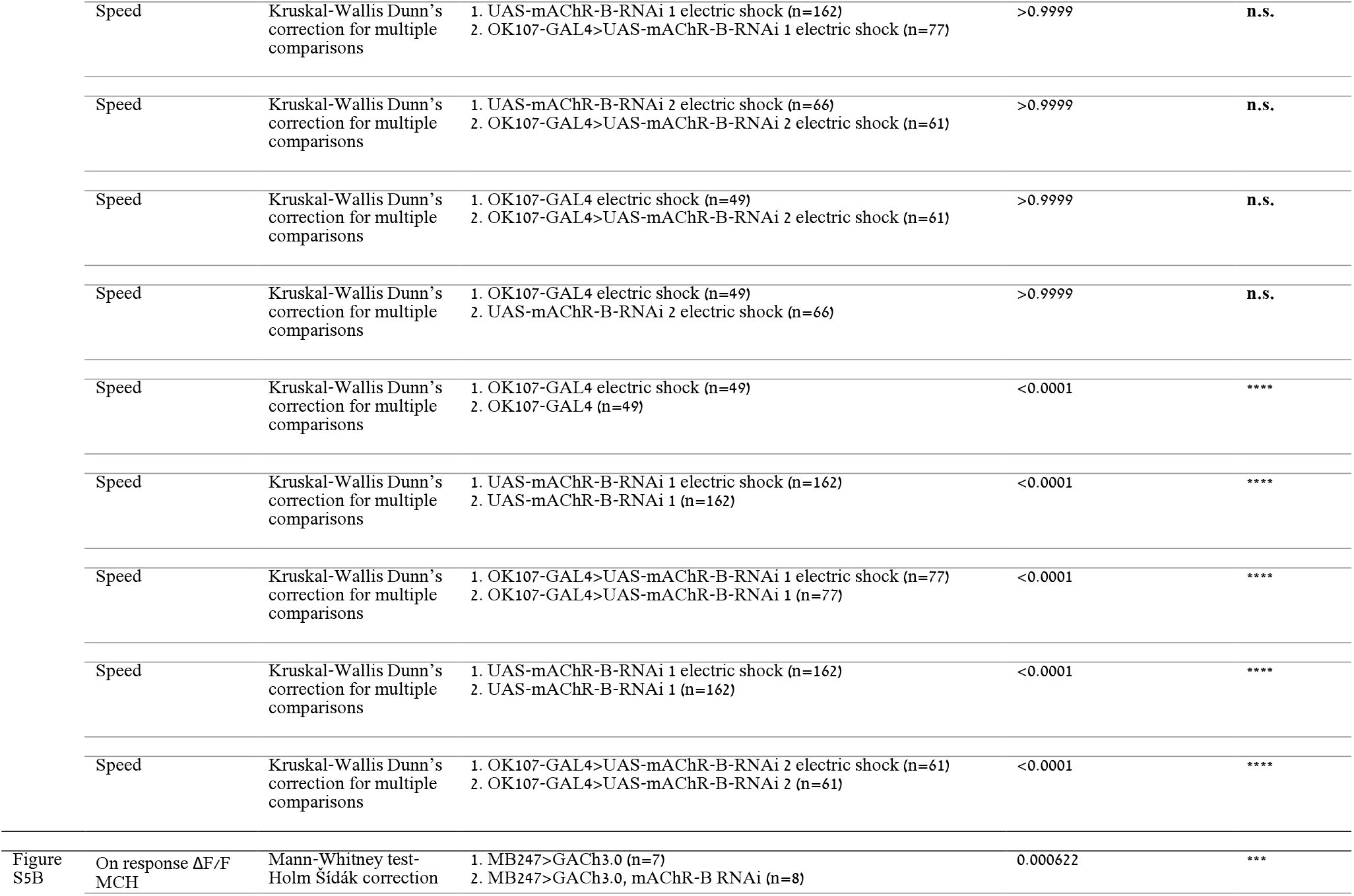
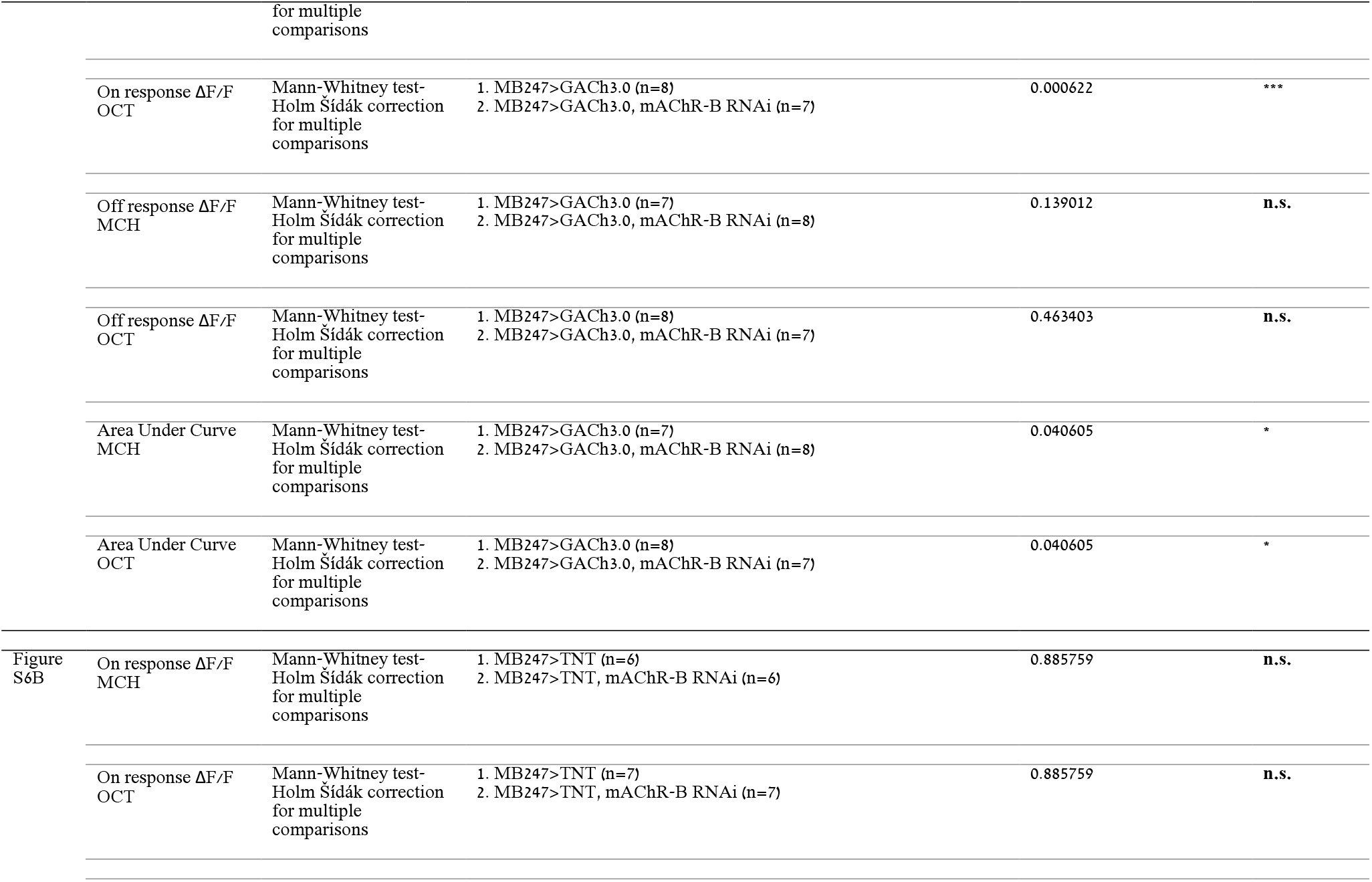
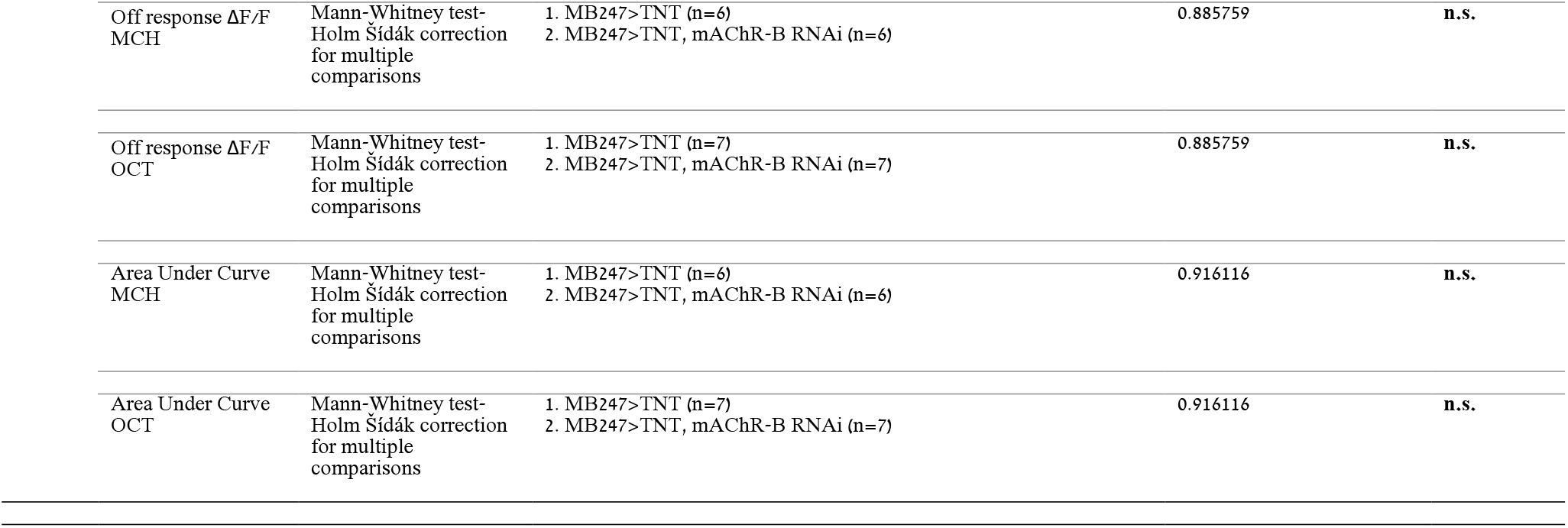
statistical analysis.

